# The RNA-binding protein DRBD18 regulates processing and export of the mRNA encoding *Trypanosoma brucei* RNA-binding protein 10

**DOI:** 10.1101/2021.09.13.460056

**Authors:** Tania Bishola Tshitenge, Bin Liu, Christine Clayton

## Abstract

The parasite *Trypanosoma brucei* grows as bloodstream forms in mammalian hosts, and as procyclic forms in tsetse flies. Trypanosome protein coding genes are arranged in polycistronic transcription units, so gene expression regulation depends heavily on post-transcriptional mechanisms. The essential RNA-binding protein RBP10 is expressed only in mammalian-infective forms, where it targets procyclic-specific mRNAs for destruction. We show that developmental regulation of RBP10 expression is mediated by the exceptionally long 7.3 Kb 3’-UTR of its mRNA. Different regulatory sequences that can independently enhance mRNA stability and translation in bloodstream forms, or destabilize and repress translation in procyclic forms, are scattered throughout the 3’-UTR. The RNA-binding protein DRBD18 is implicated in the export of a subset of mRNAs from the nucleus in procyclic forms. We confirmed that in bloodstream forms, DRBD18 copurifies the outer ring of the nuclear pore, mRNA export proteins and exon junction complex proteins. Loss of DRBD18 in bloodstream forms caused accumulation of several shortened *RBP10* mRNA isoforms, with loss of longer species, but RNAi targeting the essential export factor MEX67 did not cause such changes, demonstrating specificity. Long *RBP10* mRNAs accumulated in the nucleus, while shorter ones reached the cytoplasm. We suggest that DRBD18 binds to processing signals in the *RBP10* 3’-UTR, simultaneously preventing their use and recruiting mRNA export factors. DRBD18 depletion caused truncation of the 3’-UTRs of more than 100 other mRNAs, suggesting that it has an important role in regulating use of alternative processing sites.

## Introduction

Kinetoplastids are unicellular flagellated parasites that infect mammals and plants. The African trypanosome *Trypanosoma brucei* is a Kinetoplastid that causes sleeping sickness in humans in Africa and infects livestock throughout the tropics, with a substantial economic impact (1). *T. brucei* are transmitted by a definitive host, the Tsetse fly, or during passive blood transfer by biting flies. The parasites multiply extracellularly as long slender bloodstream forms in the mammalian blood and tissue fluids, escaping the host immune response through antigenic variation of the Variant Surface Glycoproteins (VSGs) (2). High cell density triggers growth arrest and a quorum sensing response, prompting differentiation of the long slender bloodstream forms to stumpy forms (3). The stumpy form is pre-adapted for differentiation into the procyclic form, which multiplies in the Tsetse midgut, although stumpy differentiation is not essential for Tsetse infection (4). The transition from the mammalian host to the Tsetse fly entails a decrease in temperature, from 37°C to between 20 and 32°C; and a switch from glucose to amino acids as the main source of energy, with a concomitant change to dependence on mitochondrial metabolism for energy generation (5,6). Meanwhile, the VSG coat is replaced by the procyclins (7,8). Procyclic forms later undergo further differentiation steps before developing into epimastigotes and then mammalian-infective metacyclic forms in the salivary glands (9). The developmental transitions in the *T. brucei* life cycle are marked by extensive changes in mRNA and protein levels (10-18). Differentiation of trypanosomes from the bloodstream forms to the procyclic form can be achieved in vitro through addition of 6 mM cis-aconitate and a temperature switch from 37°C to 27°C, followed by a medium change.

In Kinetoplastids, nearly all protein-coding genes are arranged in polycistronic transcription units. Mature mRNAs are generated from the primary transcript by 5’-*trans*-splicing of a 39nt capped leader sequence, and by 3’-polyadenylation (19,20). The parasite regulates mRNAs mainly by post-transcriptional mechanisms, supplemented, in the case of some constitutively abundant mRNAs, by the presence of multiple gene copies. Regulation of mRNA processing, degradation, and translation are therefore central to parasite homeostasis, and for changes in gene expression during differentiation (15-18). The sequences required for regulation of mRNA stability and translation often lie in the 3’-UTRs of the mRNAs, and most regulation so far has been found to depend on RNA-binding proteins (21). Examples include PUF9, which stabilises S-phase mRNAs (22); ZC3H11, which stabilises chaperone mRNAs (23); RBP6, which promotes differentiation of procyclic forms to epimastigotes (24,25); two PuREBPs, which are implicated in purine-dependent repression of a small number of mRNAs (26); and CFB2, which is required for stabilization of the *VSG* mRNA in bloodstream-form trypanosomes (27). RBP10 (Tb927.8.2780), which is exclusively expressed in bloodstream forms and metacyclic forms (10,11,28-30) specifically associates with procyclic-specific mRNAs that contain the motif UA(U)_6_ in their 3’-UTRs, targeting them for destruction (31). Depletion of RBP10 from bloodstream forms gives cells that can only grow as procyclic forms (31) and RBP10 expression in procyclic forms makes them able to grow only as bloodstream forms (31,32). Correct developmental regulation of RBP10 is therefore critical throughout the parasite life cycle.

In comparison with mRNA decay and translation, our knowledge of regulatory events during mRNA processing, and especially mRNA export, is rather limited. The mechanism of mRNA *trans*-splicing is basically similar to that for *cis* and *trans* splicing in other eukaryotes (20,33), as is the polyadenylation machinery (34). Unusually, however, the two processes are inextricably linked: the polyadenylation site of each mRNA is determined solely by the location of the next downstream splice site (35-38); and chemical or RNAi-mediated inhibition of either splicing or polyadenylation stops both processes (39-43). Bioinformatic analysis and limited reporter experiments indicate that - as in other eukaryotes - a polypyrimidine tract serves as a *trans*-splicing signal (12,36,37,44,45), but in contrast with at least some eukaryotes (46), there is no consensus branch-point sequence. The results of reporter experiments indicate that the nature of the sequence downstream of the splice site, as well as the length and composition of the polypyrimidine tract, influence splice site choice (47-49). There are also indications that some mRNAs are more efficiently spliced than others (16,17,50). This could only affect mRNA abundance if processing competes with degradation and indeed, it has been shown that, at least when processing is inhibited, the precursors are substrates for the exosome (51). Several RNA-binding proteins have been shown to influence splicing patterns, (52-54) but their mechanisms of action are not yet clear; often, they have double functions, also binding to the 3’-UTRs of mature mRNAs and affecting mRNA stability.

The inner ring of the trypanosome nuclear pore is very similar to those of Opisthokonts, but outer components are more diverged, giving the pore a symmetrical structure (55). Trypanosomes contain the major export factor MEX67 (Tb927.11.2370) (56,57), the helicase MTR2/p15 (Tb927.7.5760) and a homologue of Gle2/Rae1 (Tb927.9.3760). The MEX67-MTR2 complex also includes a transportin-like protein, IMP1 (Tb927.9.13520), which is required for mRNA export (57). Since export of mRNAs can be initiated before they have been completely transcribed, recognition must depend on the 5’-end: we do not know whether the export machinery recognises the cap, cap-binding proteins, or the exon junction complex. The double RNA-binding protein 18 (DRBD18) was initially identified as a substrate of arginine methylation (58). It is expressed in both bloodstream and procyclic forms. DRBD18 was shown to interact with MTR2 and promote the export of a subset of mRNAs in procyclic forms. Loss of DRBD18 caused partial nuclear retention of Mtr2 and Mex67 and showed different effects on cytoplasmic and whole cell mRNA populations (59).

Like other eukaryotes, trypanosomes have an exon junction complex, which is presumably deposited at the end of the spliced leader after or during processing: it includes Y14, Magoh and an NTF2-like protein (Tb927.10.2240) (60). Specific association of the putative eIF4AIII homologue, Tb927.11.8770 (61) with the exon junction complex has not been demonstrated, but it is mainly in the nucleus. There is also evidence that the protein from the related parasite *Trypanosoma cruzi* shuttles between nucleus and cytoplasm, and that its export depends on MEX67 (62).

In this paper, we initially aimed to unravel the mechanisms that are responsible for the stage specific expression of RBP10. We found that sequences mediating the stage specific expression of RBP10 are scattered throughout the 7.3 kb 3’-UTR of its mRNA, and that an AU repeat stimulated expression. DRBD18 depletion led to loss of RBP10 protein, despite accumulation of a truncated *RBP10* mRNA. Loss of DRBD18 led to altered processing of *RBP10* and over 100 other mRNAs. These results, combined with strong association of DRBD18 with the outer ring of the nuclear pore, implicate DRBD18 in control of mRNA processing as well as export.

## Results

### The *RBP10* 3’-UTR is sufficient for developmental regulation

Our first experiments were designed to check high-throughput results and obtain a more detailed picture of RBP10 regulation. Results from RNA-Seq and ribosome profiling (18) strongly suggest that the 3’-UTR is about 7.3 kb long, giving a total mRNA length of about 8.5 kb (Figure 1A). The middle region of the 3’-UTR is present in one other contiguous sequence in the TREU927 genome assembly (giving the grey-coloured reads in the alignment in Figure 1A) but this sequence is found only once in the Lister 427 strain genome (63). In our experiments, we used two *T. brucei* strains: either Lister 427 which cannot complete differentiation, or EATRO1125, which is differentiation-competent but for which an assembled genome is not yet available. The gene immediately downstream of *RBP10* (Tb927.8.2790) is annotated as an acetyl-coA synthetase pseudogene. A complete acetyl-coA synthetase coding region is present elsewhere in the genome, but the presence of read alignments over the region that surrounds the Tb927.8.2790 coding region suggests that the pseudogene mRNA is also present in both bloodstream and procyclic forms.

**Figure 1.**
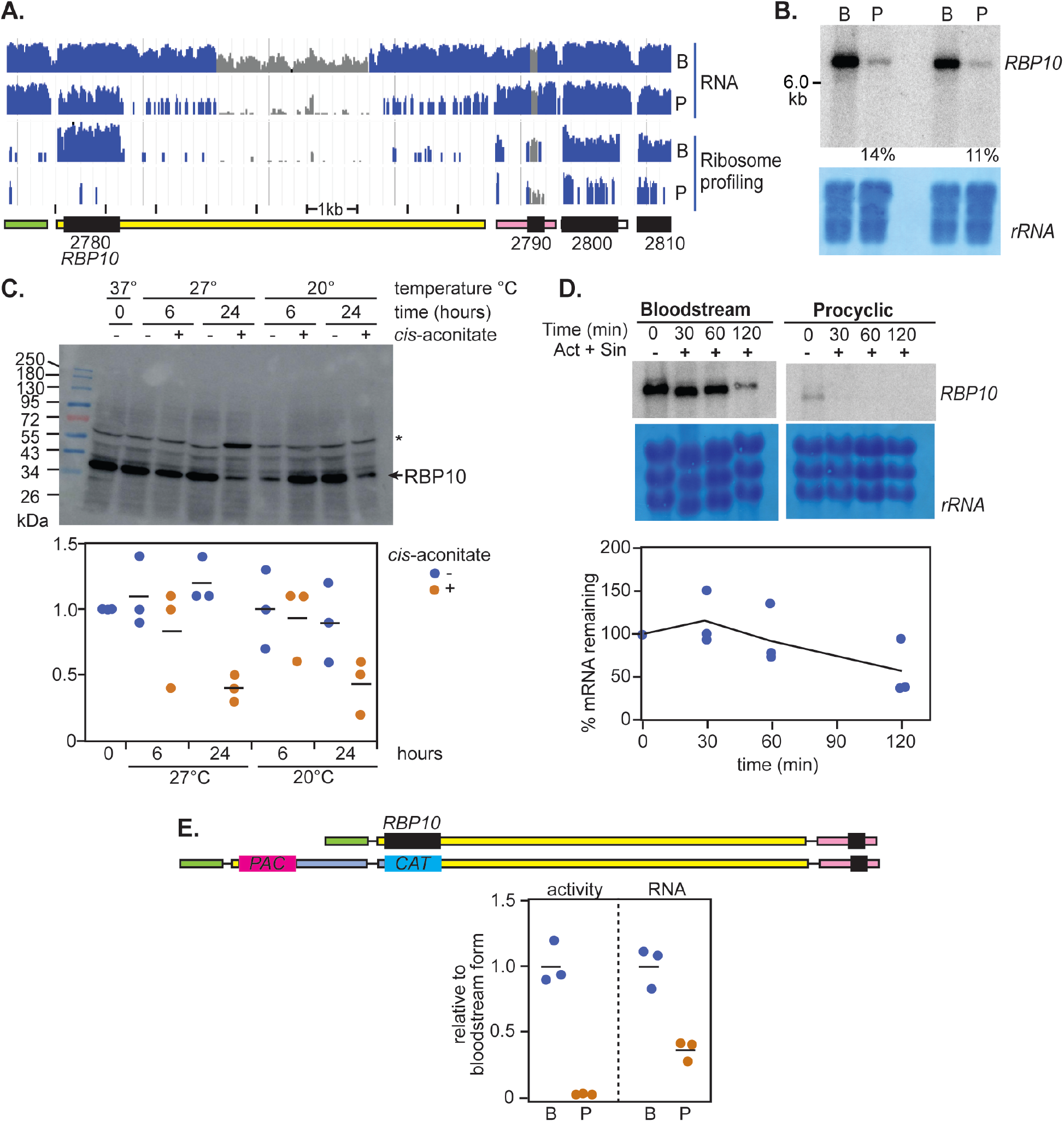
Developmental regulation of the *RBP10* mRNA. **A**. RNA-sequencing data and ribosome profiling (15) reads for bloodstream forms (B) and procyclic forms (P), aligned to the relevant segment of the TREU927 reference genome. Unique reads are in blue and non-unique are in grey, on a log scale. The sequences in the *RBP10* (Tb927.8.2780) 3’-UTR designated as non-unique in this image are also present in an TREU927 DNA contiguous sequence that has not been assigned to a specific chromosome; this segment is absent in the Lister427 (2018) genome. The positions of open reading frames (black) and untranslated regions have been re-drawn, with the initial “Tb927.8” removed for simplicity. Transcription is from left to right. Data from all remaining panels are for the EATRO1125 strain. **B**. Northern blot for two independent cultures showing the relative mRNA abundance of *RBP10* mRNA in bloodstream forms and in procyclic forms. A section of methylene blue staining is depicted to show the loading and the measured amounts in procyclic forms, relative to bloodstream forms, are also shown. **C**. Regulation of RBP10 protein expression during differentiation. Cells were incubated with or without 6 mM *cis*-aconitate at the temperatures and for the times indicated. The asterisk shows a band that cross-reacts with the antibody. **D**. Half-life of *RBP10* mRNA. Cells were incubated with Actinomycin D (10 µg/ml) and Sinefungin (2 µg/ml) to stop both mRNA processing and transcription. The amount of *RBP10* mRNA in bloodstream forms was measured in three independent replicates. The initial delay or even increase in abundance is commonly seen for relatively stable trypanosome mRNAs, and is of unknown origin. No *RBP10* mRNA was detectable after 30min similar treatment of procyclic forms. **E**. The *RBP10* 3’-UTR is sufficient for regulation. A dicistronic construct mediating puromycin resistance (*PAC* gene) and encoding chloramphenicol acetyltransferase (*CAT* gene) replaced one of the two *RBP10* open reading frames (upper panel) in bloodstream forms. These then were differentiated to procyclic forms. The lower panel shows measurements of CAT activity and mRNA for three independent cell lines, normalised to the average for bloodstream forms.

Previous transcriptome results indicated that there are about 4 copies of *RBP10* mRNA per cell, and slightly under 1 per cell in procyclics, but that the ribosome density on the coding region is 9 times higher in bloodstream forms (16,17). Northern blot results for the EATRO1125 strain showed an *RBP10* mRNA that migrated slower than the 6kb marker (Figure 1B). Extrapolation suggested a length of about 8.5 kb. In this experiment, there was 8-fold more *RBP10* mRNA in bloodstream forms than in procyclic forms (Figure 1B). The amount of RBP10 decreased about 3-fold after 24h incubation with 6mM *cis*-aconitate at either 20°C or 27°C, and both *cis*-aconitate and the temperature drop were required for the regulation (Figure 1C). We had previously measured, by RNA-Seq, an *RBP10* mRNA half-life of just over 1h in Lister 427 bloodstream forms (16), while a new individual measurement in EATRO1125 suggested a half-life of 2h (Figure 1D). For procyclic forms, the RNA-Seq replicates were very poor (16), but from Northern blotting the half-life was probably less than 30 min (Figure 1D). Finally, to find out whether the 3’-UTR was sufficient for regulation, we integrated a chloramphenicol acetyl transferase (*CAT*) gene into the genome of strain EATRO1125 bloodstream forms, directly replacing one *RBP10* allele (Figure 1E). After differentiation to procyclic forms, *CAT* mRNA was about 3-fold down-regulated, but there was no detectable CAT activity (Figure 1E). This shows that the *RBP10* 3’-UTR is sufficient for developmental regulation.

### The *RBP10* 3’-UTR contains numerous regulatory elements

In order to dissect regulatory elements inside the *RBP10* 3’-UTR that contribute to the stability and translation of *RBP10* mRNA in bloodstream forms, or to its instability and translational repression in procyclic forms, we made use of a reporter plasmid that integrates into the tubulin locus, resulting in read-through transcription by RNA polymerase II. Cell lines can be selected with G418 using the *NPT* (neomycin phosphotransferase) marker. The *CAT* reporter mRNA has a 5’-UTR and splice signal from an *EP* procyclin gene (Figure 2A). At the 3’-end between *CAT* and *NPT* is an intergenic region from between the two actin (*ACT*) genes, with a restriction site exactly at the mapped polyadenylation site (Figure 2A). The reporter produces in a *CAT* mRNA bearing an *ACT* 3’-UTR, with polyadenylation driven by the downstream splice site for *NPT*. In all experiments, the reporter plasmid was transfected into EATRO1125 bloodstream forms and two or three independent clones were then differentiated into procyclic forms. CAT activities were measured enzymatically and mRNA levels were measured by Northern blotting, which simultaneously allowed us to check the sizes of the mRNAs. All values were normalised to arithmetic mean results from the *ACT* 3’-UTR control. The sizes of the mRNAs are tabulated in Supplementary Table S1 and the sequences are in Supplementary text 1. Most of the reporter mRNAs migrated either as expected or slightly faster. The latter suggests polyadenylation upstream of the expected sites.

**Figure 2.**
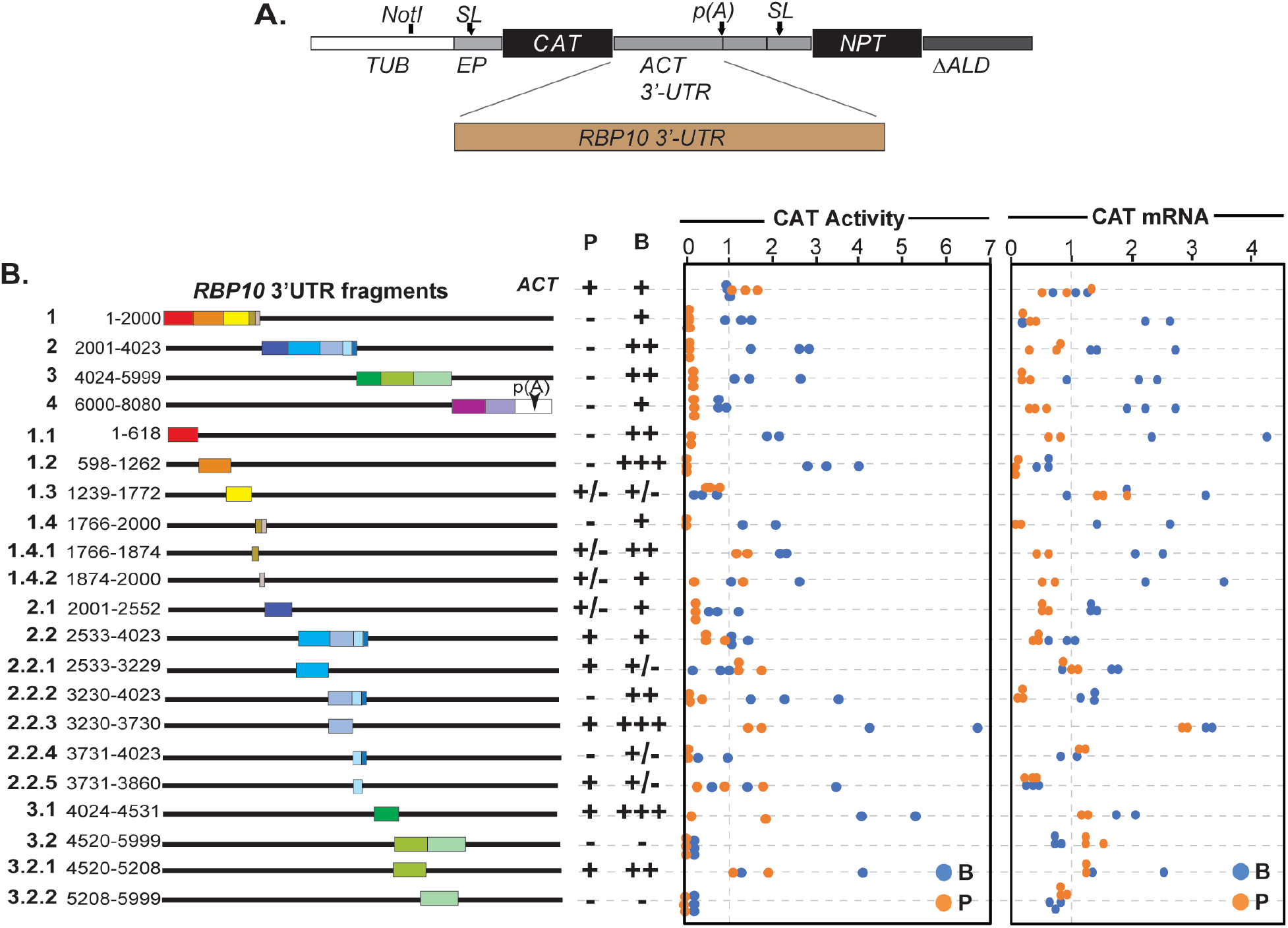
The *RBP10* 3’-UTR contains several regulatory sequences. **A**. Cartoon showing the *CAT* reporter construct (pHD2164) used in this study. The different fragments of the *RBP10* 3’-UTR were cloned downstream of the *CAT* reporter coding sequence. The polyadenylation site “p(A)” is specified by the *NPT* splice signal. *SL* indicates the spliced leader addition sites. *ACT* denotes actin. **B**. Schematic diagram of the *RBP10* 3’-UTR segments used to map the regulatory sequences. Measurements are on the right. Each dot is a result for an independent clone, with blue and orange representing bloodstream (B) and procyclic (P) forms, respectively. Values were normalized to those for the *ACT* 3’-UTR control (pHD2164). The average result for the control BSF cell line was set to 1. For “expression”, “+” = 0.5-2x, “++”=2-3x and “+++” >3x.

First, we examined four different fragments (1-4 in Figure 2B), each roughly 2 Kb long. Fragment 4 extends beyond the RBP10 polyadenylation site, including the intergenic region before downstream gene (Tb927.8.2790); and the size of the resulting RNA suggested use of the genomic processing signals (Supplementary Table S1). To our surprise, all four fragments decreased *CAT* mRNA, and abolished CAT activity in procyclic forms; fragments 2 and 3 also gave enhanced CAT activity in bloodstream forms (Figure 2B). Clearly several regulatory sequences were present. Further dissection showed that fragments 1.1, 1.2, 1.4 and 2.2.2 (Figure 2B, C) were each able to suppress the CAT reporter levels in procyclic forms and increase the expression levels in bloodstream forms. Fragment 1.4 (Figure 2C) was the shortest that we obtained that was able to decrease expression in procyclic forms. Fragments 1.3, 3.2 and 3.2.2 gave low CAT expression levels in both stages. It was notable also that sometimes, fragmentation of a sequence created activity that had been absent previously: for example, fragment 2.2 gave poor regulation but the sub-fragment 2.2.2 regulated like fragment 2, and 2.2.3 gave higher activity in bloodstream forms, approaching that of 3.1 (Figure 2B). These results show that regulation of RBP10 expression is achieved by numerous sequence elements, some of which may have competing activities.

To look for regulatory motifs, we compared different sets of sequences using MEME or DREME (64,65). We first searched for motifs enriched in the *RBP10* regulatory fragments relative to the those that lacked regulation, and then for motifs that were enriched in both the regulatory fragments and other co-regulated bloodstream-form specific mRNAs. Other non-regulated or procyclic-specific mRNAs were used as controls. We found no enriched motifs, suggesting that several different sequences (and perhaps RNA-binding proteins) are implicated in the regulation.

### AU-Rich elements affect expression

We next examined two repetitive motifs. Deletion of the (AU)_10_ element from *RBP10* 3’-UTR fragment 1.2 resulted in a drastic decrease of reporter expression levels (Figure 3A). Several fragments of the *RBP10* 3’-UTR with good translation in bloodstream forms contain poly(A) tracts, and we therefore speculated that they might act by recruiting a poly(A) binding protein. However, deletion of the poly(A) tracts from one of these fragments unexpectedly resulted in an increase in reporter expression levels, rather than a decrease (Figure 3B). Therefore, so far, the only short motif demonstrated to enhance expression was the (AU) repeat. We do not know whether this is stage-specific or not.

**Figure 3.**
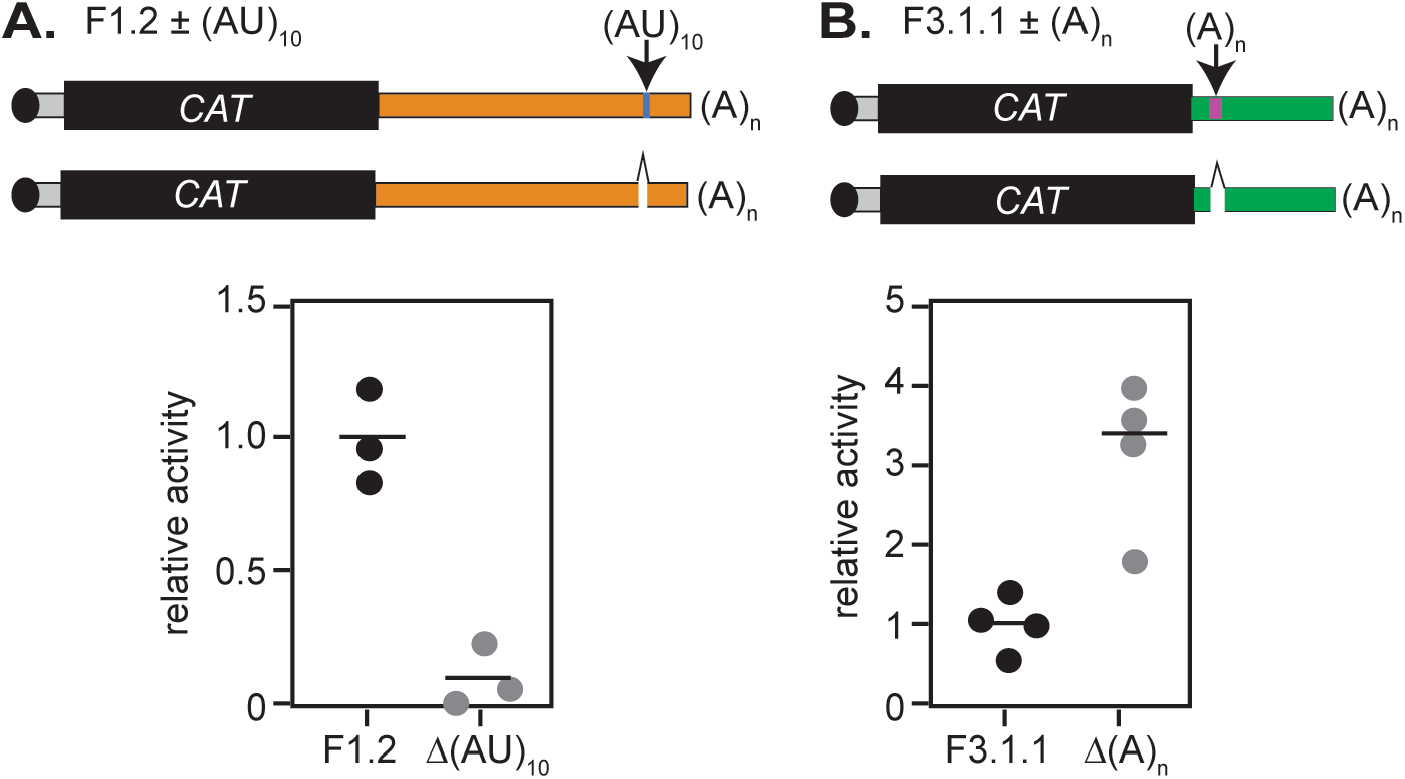
Analysis of specific sequences. **A**. Results for a reporter plasmid containing *RBP10* 3’-UTR fragment 1.2 bearing (AU)_10_ repeats and a mutated version without them. The average result for the fragment with the repeats is set to 1. **B**. As (A) but showing deletion of a poly(A) tract from fragment 3.1.1.

### DRBD18 depletion affects *RBP10* mRNA processing

To investigate which RNA-binding proteins might affect RBP10 expression, we looked for two types of profile. For enhancers, these would be proteins that are expressed (and perhaps essential) in bloodstream forms, can bind mRNA in bloodstream forms and can enhance expression when tethered to a reporter mRNA. For repressors, we looked for proteins that are expressed (and perhaps essential) in procyclic forms, can bind mRNA and can repress expression when tethered. We also examined the existing literature on such proteins, looking for those that bound to and increased *RBP10* mRNA in procyclic forms or decreased it in bloodstream forms. From this survey, selected proteins were depleted by RNAi and the effects on RBP10 protein expression were assayed (Supplementary Table S2). A candidate procyclic-specific repressor, ZC3H22, was previously studied and not implicated in *RBP10* control (66). We found that RNAi targeting HNRNPF/H, ZC3H28, ZC3H40 or ZC3H45 in bloodstream forms had no effect on RBP10 expression. One remaining candidate, ZC3H44, was not investigated.

We selected DRBD18 for study because its depletion in procyclics was shown to increase *RBP10* mRNA levels (58). To find out whether this also resulted in an increase in RBP10 protein, we made EATRO1125 bloodstream forms with inducible RNAi targeting DRBD18. Depletion of DRBD18 in bloodstream forms caused cell growth arrest after 24 hours (Supplementary Figure S1A). The concomitant decrease of RBP10 protein (Supplementary Figure S1B) could be a direct effect, or caused by slower growth. The same cells were differentiated to procyclic forms. Now, DRBD18 depletion was incomplete and transient; but after 2 days, a trace of RBP10 expression was detectable (Supplementary Figure S1C).

We also checked the location of DRBD18, since existing results were contradictory: (58) reported that it was in the cytosol whereas C-terminally GFP-tagged DRBD18 was in the nucleus (67). The control cytoplasmic protein, RBP10, and the nuclear exoribonuclease XRND (68) showed the expected distributions. In contrast, we found that DRBD18 was associated with both compartments (Supplementary Figure S1D, E), which is consistent with a role in nuclear export.

Since the effects of DRBD18 depletion in procyclic forms had been analysed extensively elsewhere, we decided to focus on bloodstream forms. We decided to use Lister 427 strain trypanosomes for these experiments, because they grow to higher densities than EATRO1125. DRBD18 depletion caused clear growth inhibition (Figure 4A) and a decrease in RBP10 protein (Figure 4B). Remarkably, DRBD18 depletion also caused accumulation of progressively smaller *RBP10* mRNAs (Figure 4C); since the blot was hybridised to a coding region probe, these must be products of alternative polyadenylation. To find out whether DRBD18 depletion generally inhibits processing, we examined mRNA from the tandemly repeated tubulin genes, because processing inhibition causes accumulation of tubulin RNA dimers and multimers. Notably, processing of *β-tubulin* mRNA was not affected, showing that the effect on *RBP10* mRNA was specific (Figure 4C). A similar effect on *RBP10* mRNA was seen in DRBD18-depleted EATRO1125 strain bloodstream forms (Supplementary Figure S1F, G).

**Figure 4:**
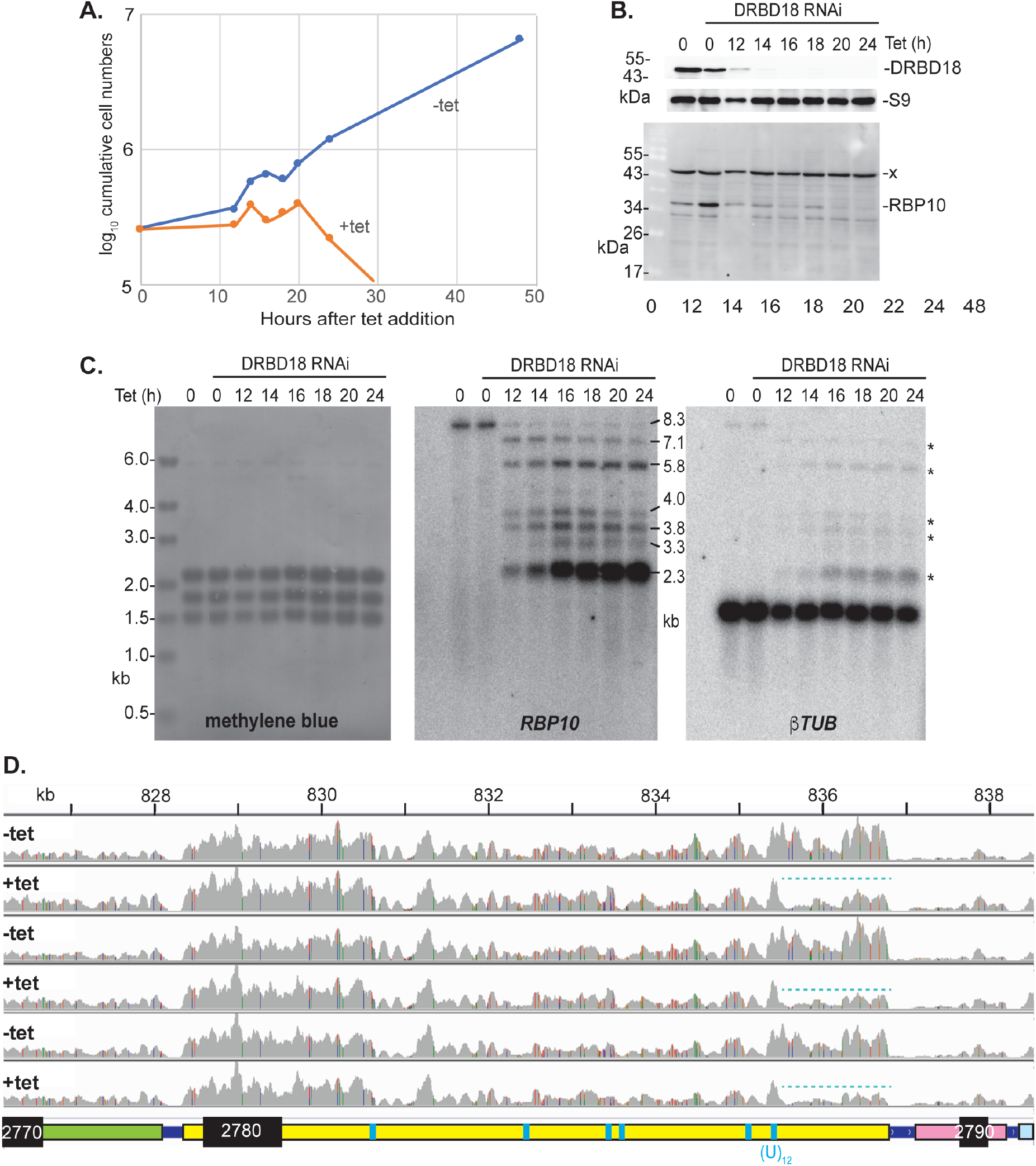
Effect of *DRBD18* RNAi on *RBP10* mRNA and expression levels. **A**. Example of cumulative growth curves of bloodstream forms with and without induction of *DRBD18* RNAi. **B**. DRBD18 and RBP10 protein levels after RNAi induction, corresponding to (A). Ribosomal protein S9 is the loading control. Anti-RBP10 antibodies (28) were used for detection of RBP10 and a non-specific band (x). **C**. RNA from the samples used in (A) was used to examine effects on *RBP10* mRNA. The methylene blue-stained membrane is shown on the left, with *RBP10* hybridisation in the centre. The membrane was then stripped and hybridised with a beta-tubulin probe as a further control; \**RBP10* signal that remained after stripping. Analysis of 3’-ends of the different *RBP10* mRNAs by RT-PCR (3’-RACE) using oligo (dT)_18_ was not successful, perhaps because the sequence is generally of low complexity and the forward primers chosen were similar to sequences in other mRNAs. **D**. Visualisation of RNA-Seq results after 12h tetracycline-mediated induction of *DRBD18* RNAi, using the Integrated Genomics Viewer (69,70). Three replicates are shown. Note that these results are shown on a linear scale whereas those in Figure 1A are on a log scale. Proposed full length mRNAs are shown below the tracks; positions of (U)_12_ or (U)_6_C(U)_6_ sequences are indicated in cyan but there are numerous other polypyrimidine tracts present. The cyan dotted line highlights loss of reads at the end of the *RBP10* 3’-UTR after 12h *DRBD18* RNAi. The read densities over the middle portion of the 3’-UTR are lower than the rest, probably because some of the reads will be aligned to the additional sequence copy in the TREU927 reference genome. Coloured lines indicate read mismatches with the TREU927 reference genome.

To confirm these results, we examined the transcriptomes of bloodstream forms 12 hours after DRBD18 RNAi induction. This time was chosen as the first point at which DRBD18 was completely depleted (Figure 4B) without much effect on growth (Figure 4A). Overall, the numbers of *RBP10* reads were not much affected at this time point (Supplementary Table S3). Visualisation in the Integrated Genome Viewer (69,70) confirmed that with or without tetracycline induction of RNAi, the read density over the RBP10 coding region was 2-3 times higher than the density over the upstream gene, Tb927.8.2770 (Figure 4D). However, over the last 2kb of the 3’-UTR, there were far fewer reads after DRBD18 depletion (cyan dotted line), indicating loss of the longest *RBP10* mRNAs and consistent with the Northern blot result.

The results so far suggested that multiple alternative polyadenylation sites were being used after *DRBD18* depletion. We were unable to find out whether the downstream mRNA, from the acetyl-coA synthetase pseudogene, was correspondingly abnormally spliced, because probes for the coding region gave only bands that migrated slower than expected - presumably because they also hybridize with mRNA from elsewhere in the genome that encodes the full-length protein. We tried using a non-coding probe from the intergenic region but the RNAs detected were again much larger than expected and probably of non-specific origin.

A previous paper has implicated DRBD18 in export of mRNAs from the nucleus. We therefore looked to see whether inhibition of nuclear export by depletion of MEX67 also caused altered processing of *RBP10* mRNA. MEX67 is required for mRNA export and its depletion causes general accumulation of mRNAs in the nucleus (56). After 24h RNAi induction, growth of the parasites was clearly inhibited (Figure 5A) and *MEX67* mRNA was reduced (Figure 5B), but the migration of *RBP10* mRNA was unaffected (Figure 5B). The altered processing of *RBP10* mRNA after DRBD18 depletion is therefore not caused only by retention in the nucleus.

**Figure 5:**
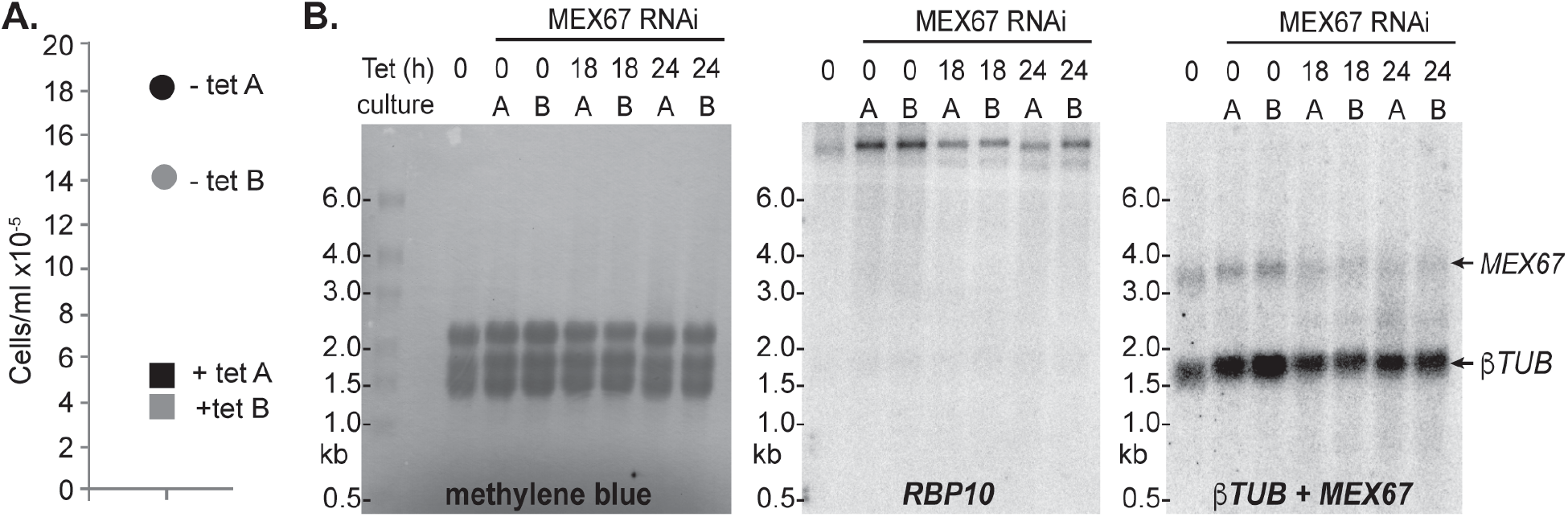
Effect of *MEX67* RNAi *on RBP10* mRNA and expression levels. **A**. Cell densities in two cultures 24h after addition of tetracycline to induce *MEX67* RNAi. **B**. Northern blots using RNA from the two cultures, with detection of rRNA (methylene blue) and of *RBP10* and beta-tubulin mRNAs.

### Interactions of DRBD18

DRBD18 interacts directly with MTR2, part of the complex required for export of mRNAs from the nucleus, and also pulls down MEX67 from procyclic form extracts (59). Non-quantitative mass spectrometry results with DRBD18 preparations from procyclic forms also suggested co-purification of some nuclear pore components and various RNA-binding proteins (58). Since label-free quantitative mass spectrometry is now readily available, we used it to re-examine DRBD18 protein associations in more detail. DRBD18 with a C-terminal TAP (tandem affinity purification) tag was expressed from one allele in bloodstream forms. We do not know whether this protein was fully functional, since attempts to delete the other (untagged) copy failed. It was not possible to obtain cells expressing DRBD18 with N-terminal tags.

We used just a single step of purification in order to avoid loss of material and interaction partners. DRBD18-TAP was pulled-down using IgG beads, then released by Tobacco Etch Virus (TEV) protease cleavage at a site within the tag. The resulting mixture was analysed by quantitative mass spectrometry. GFP-TAP and TAP-ZC3H28 (71) served as controls. ZC3H28 is an RNA-binding protein that is predominantly in the cytoplasm (71).

Figure 6 shows analysis of the data using the Perseus algorithm (72). The major advantage of Perseus is that it yields probability values that are adjusted for multiple testing. A disadvantage is that it handles absent values by substituting simulated intensities, which can lead to false negative results if a protein that interacts with the investigated protein is not detected at all in the negative control. In the following discussion, we consider two overlapping sets of proteins as being significantly enriched: those that had adjusted P-values of less than 0.01, with greater than 4-fold enrichment as calculated in Perseus, and those that were detected with at least one peptide in all of the DRBD18 preparations but none of the relevant controls. Details are in Supplementary Table S4. The comparisons showed that DRBD18-TAP specifically copurified several sets of proteins. Most dramatically, there was highly specific co-purification of all components of the outer ring of the nuclear pore (55,73,74). We found MTR2 as expected, but also a transportin-2-like protein, an importin-beta subunit, a Ran binding protein, the putative nuclear RNA helicase EIF4AIII and a possible Ran-GAP. MEX67 was found in two of the three DRBD18 preparations and none of the others. The exon junction complex components Magoh and NTF2-domain-protein (60) were found with both DRBD18 and ZC3H28, but not the GFP control, though Magoh was more reproducibly associated with DRBD18. Intriguingly, DRBD18 also specifically copurified multiple components of the mitochondrial RNA editing machinery; this might warrant future investigation.

**Figure 6:**
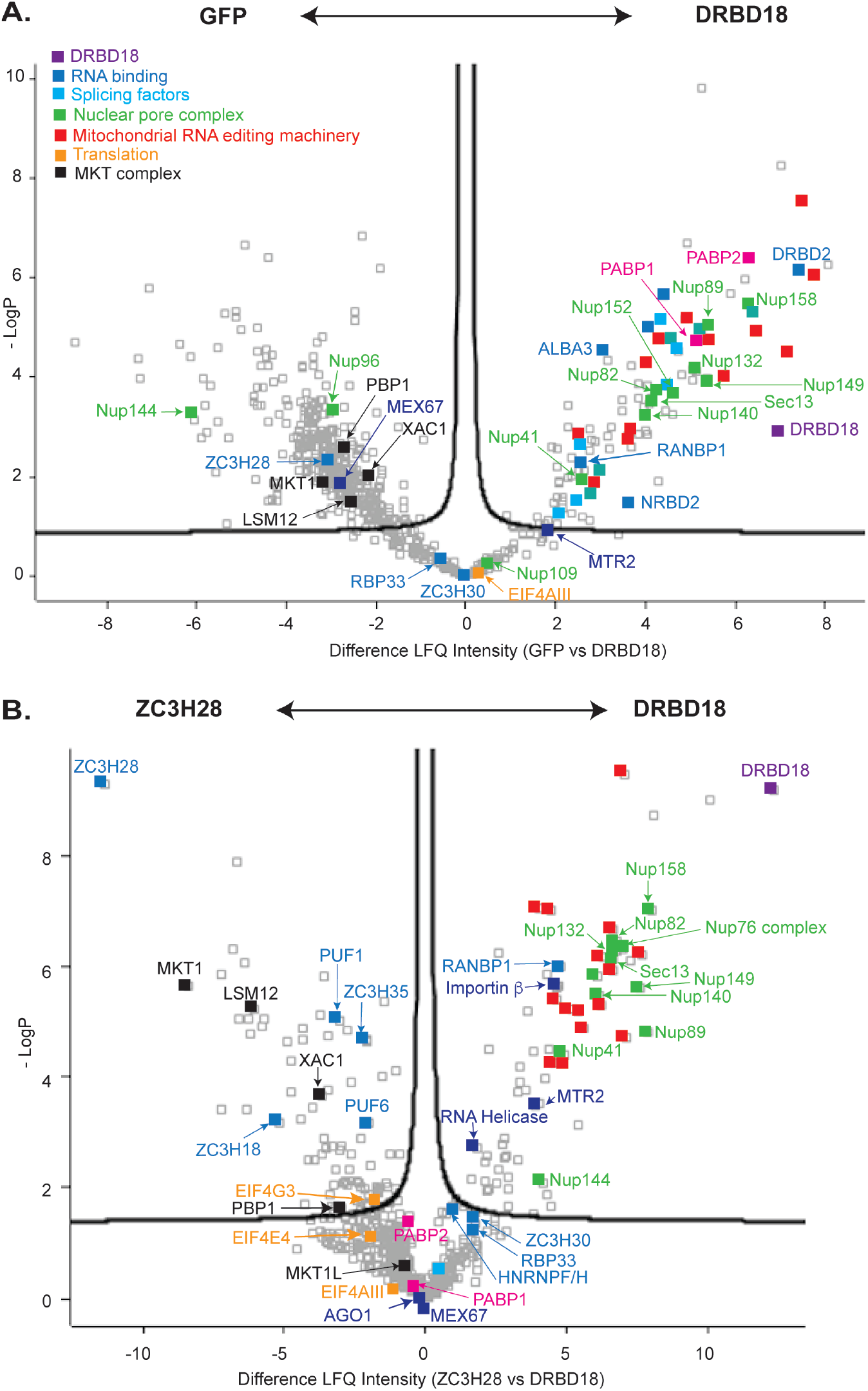
Interactions of DRBD18. **A**. Volcano plot showing proteins that were significantly enriched with DRBD18-TAP relative to TAP-GFP. Proteins of interest are labelled. The x-axis shows log_2_ enrichment, while the y-axis is -log_10_ of the P-value generated by the Perseus algorithm (72). Proteins outside the curved line are significantly enriched. **B**. Volcano plot showing proteins that were significantly enriched with DRBD18-TAP and TAP-ZC3H28. The data for ZC3H28 and GFP were described previously (71).

The RNA-binding domain proteins ZFP2, RBP33 and ZC3H30 were specifically associated with DRBD18. ZFP2 is required for differentiation of bloodstream to procyclic forms (75). The function of ZC3H30 is unknown (76). RBP33 is an essential nuclear protein, so its association with DRBD18 but not ZC3H28 corresponds to the localisations of the two proteins. RBP33 associates with non-coding RNAs, especially from antisense transcription (77), but its depletion led to loss of mRNA and spliced leader RNA (78), suggesting a link to splicing. Both DRBD18 and ZC3H28 enriched various ribosomal proteins and RNA-binding proteins, including both poly(A) binding proteins, ALBA2, ALBA3, DRBD2, DRBD3, ZC3H9, ZC3H34, ZC3H39, ZC3H40, UPF1, HNRNPH/F and TRRM1. There were also some splicing factors, the RNA interference Argonaute protein AGO1, and the translation initiation complex EIF4E3/4G4 (Supplementary table S4). Many of these proteins are likely to associate via RNA. Unlike ZC3H28, DRBD18 was not associated with MKT1, LSM12, XAC1, and PBP1, which are components of a complex that promotes translation and mRNA stability (79,80). Association of PUF1, PUF6, ZC3H35, and ZC3H18 was also specific to ZC3H28.

### Long *RBP10* transcripts are retained in the nucleus after DRBD18 depletion

Our mass spectrometry results strongly supported and expanded the previous observations implicating DRBD18 in mRNA export. Inhibition of export alone was however not sufficient to cause accumulation of shorter *RBP10* mRNA variants, and the previously publication had already indicated that the effects of DRBD18 are sequence-specific (59). We hypothesised, therefore, that DRBD18 binds to possible processing sites in the *RBP10* 3’-UTR, and that this not only promotes efficient export of the long mRNA, but also inhibits the use of the alternative sites. If this were true, then DRBD18 RNAi might selectively trap longer *RBP10* mRNAs in the nucleus.

To test the effects of DRBD18 depletion on *RBP10* mRNA localisation, we examined the locations of the different *RBP10* mRNAs by single-molecule fluorescent *in situ* hybridization. To distinguish the different mRNAs, we used four probes (1,2,3, and 4) (Figure 7A). Details of the results are in Supplementary Table S5. The shortest species detected had signals from the coding region only (pink probe 1) and were mostly nuclear: these could represent precursors that were still undergoing transcription. The next shortest mRNAs detected hybridised only to probes 1 (coding region) and 2. These must be 2-7 kb long (Figure 7A, B). The mRNAs that hybridise to probes 1 and 3 are at least of intermediate length (5-7 kb long); and those that hybridise to probes 1,2 and 4 or 1,3 and 4 are more than 7.5 kb long (Figure 7A, B). On average, 5% of the mRNAs hybridised only to probes 1 and 4: this is physically unlikely and may indicate of the background failure rate for hybridisation of probes 2 or 3 (Supplementary Table S5). The numbers of mRNAs hybridising only to probes 2 or 3 were unexpectedly high (average 17% of signals); although these do not match any other sequences in the Lister427 genome, they do contain many low complexity sequences which might result in cross-hybridisation. Very few RNAs hybridised to the green probe only; they might be degradation products (81). If all mRNAs hybridising to the coding region probe were included, wild-type cells contained 3-3.5 *RBP10* mRNAs per cell; previous calculations (16) had indeed suggested 4 *RBP10* mRNAs per cell. For the further discussion, in order to be certain that only specific signals are considered, we consider only the long, intermediate and short mRNAs (as defined above) which hybridised to at least 2 probes.

**Figure 7:**
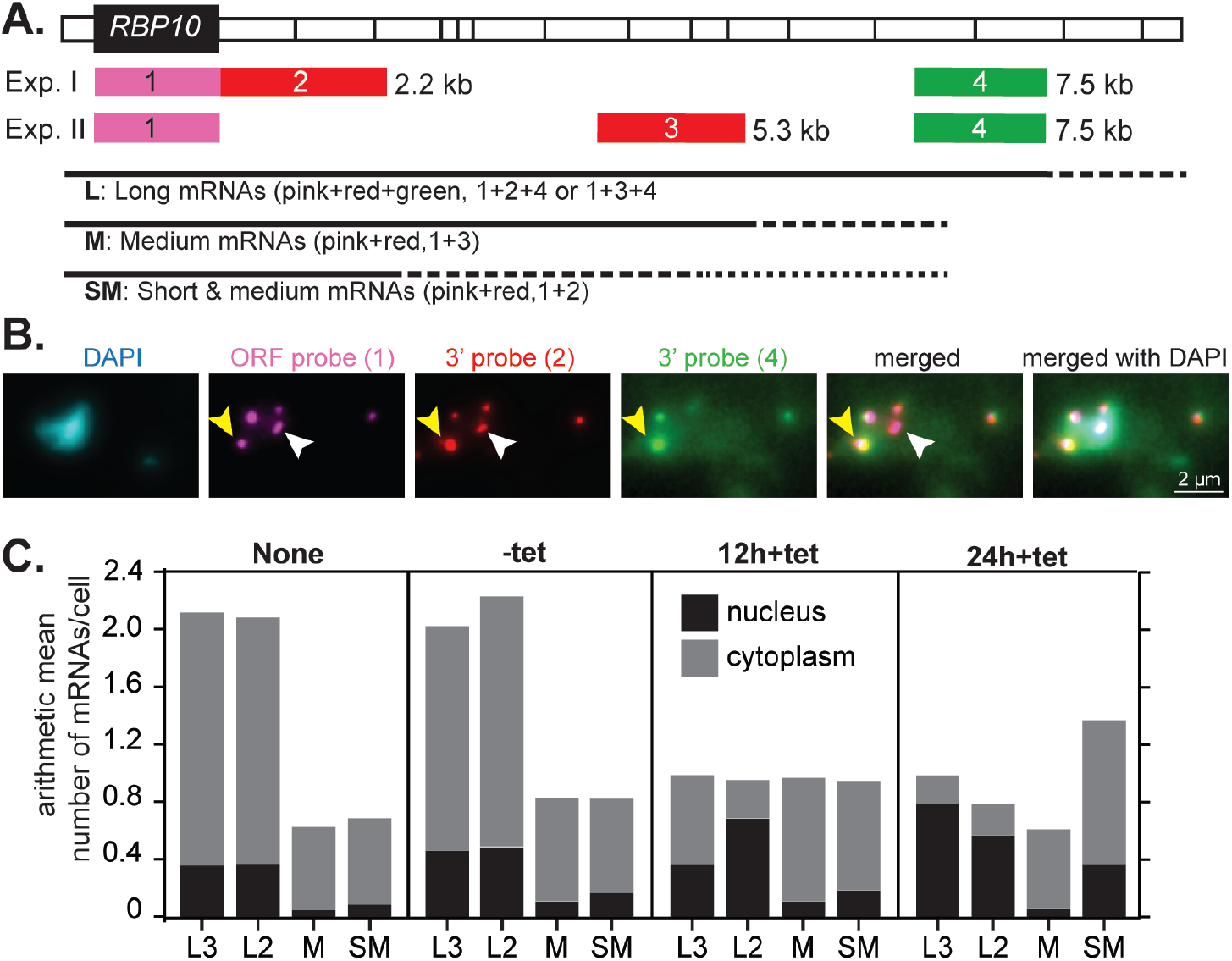
Long *RBP10* mRNAs are trapped in the nucleus after DRBD18 depletion. Individual *RBP10* mRNAs were detected by single-molecule fluorescent *in situ* hybridization with or without DRBD18 depletion for 12 or 24h. In each case, 50 cells were analysed. **A**. Map showing the positions of the probes on the *RBP10* mRNA, approximately to scale. The 3’-UTR divisions on the map correspond to those in Figure 2. The bars indicate the extent of mRNAs detected by the probe combinations. The dotted lines indicate that RNAs hybridising to the different probe sets could have various lengths. The distances next to the bars indicate the position of the probe end on the mRNA. Exp.: experiment. **B**. Example image of *RBP10* mRNAs in a wild-type cell, detected by single-molecule fluorescent *in situ* hybridization using probes 1,2 and 4 (experiment 1 in panel (A)). DNA was stained with DAPI. The white arrow indicates an mRNA that hybridises only to the coding region (open reading frame, ORF) and 3’ probe 2; the yellow arrow points to one of the four mRNAs that hybridise to probes 1,2 and 4. **C**. Mean numbers of long, medium and short mRNAs per cell under different conditions. The data were from 50 cells. “None”: Cells with no RNAi plasmid; “-tet”: RNAi cells without tetracycline; and cells after 12h or 24h tetracycline treatment. All slides were hybridised with the red, pink and green probes, and results for hybridisation with the two different red probes are shown separately. “L3” indicates long mRNAs that hybridised with probes 1,3 and 4 (experiment II) and “L2” indicates long mRNAs that hybridised with probes 1,2 and 4 (experiment I). The black bars indicate mRNAs in the nucleus and the grey bars, mRNAs in the cytoplasm. Numbers for mRNAs hybridising to only one probe, or red and green only, are in Supplementary Table S5.

In wild-type cells, about 70% of *RBP10* mRNAs were longer than 7.5 kb (Figure 7C); 17% of the long mRNAs and 9-12% of the short or medium ones were in the nucleus (Figure 7C). Cells containing the *DRBD18* RNAi plasmid, but with no induction, showed similar values to wild-type, except that the proportion of mRNAs that were in the nucleus was marginally elevated, suggesting that the RNAi was very slightly leaky (Figure 7C). 12h after RNAi induction, the total number of full-length *RBP10* mRNAs was roughly halved and most of those that remained were retained in the nucleus, especially after 24h (Figure 7C). Both the medium- and shorter *RBP10* mRNAs were reproducibly mostly exported to the cytoplasm at all time-points. Results from the experiment were consistent with those from Northern blotting: the medium-length mRNAs had increased after 12h and decreased again at 24h, while the numbers of shorter mRNAs progressively increased.

These results strongly support our hypothesis that DRBD18 is required for export of long *RBP10* mRNAs from the nucleus. In contrast, after further processing, the shorter mRNAs can be exported without DRBD18. We know that shortened *RBP10* 3’-UTRs do not prevent translation (Figure 2), so the reduced amount of RBP10 protein after DRBD18 RNAi may be caused by the decrease in total cytosolic *RBP10* mRNA (Figure 7C).

### DRBD18 depletion affects processing of at least 100 mRNAs

DRBD18 depletion affected processing of *RBP10* mRNA, but not tubulin mRNAs. But in procyclic forms, transcriptome results suggested that *DRBD18* RNAi causes retention of nearly 100 mRNAs in the nucleus (59). To find out whether *DRBD18* RNAi affects processing of additional mRNAs in bloodstream forms, we first manually examined a few mRNAs with long 3’-UTRs using the Integrative Genomics Viewer (69). Prompted by the results, we chose the *DRBD12* mRNA for further investigation by Northern blotting. Indeed, *DRBD12* mRNA - which is usually about 8.4 kb long - accumulated as a 2.6 kb mRNA from 12h *DRBD18* RNAi onwards, with an intermediate band of about 4kb (Figure 8). The genomics viewer provided clear evidence for accumulation of a 4kb mRNA (orange dotted line) after 12h.

**Figure 8.**
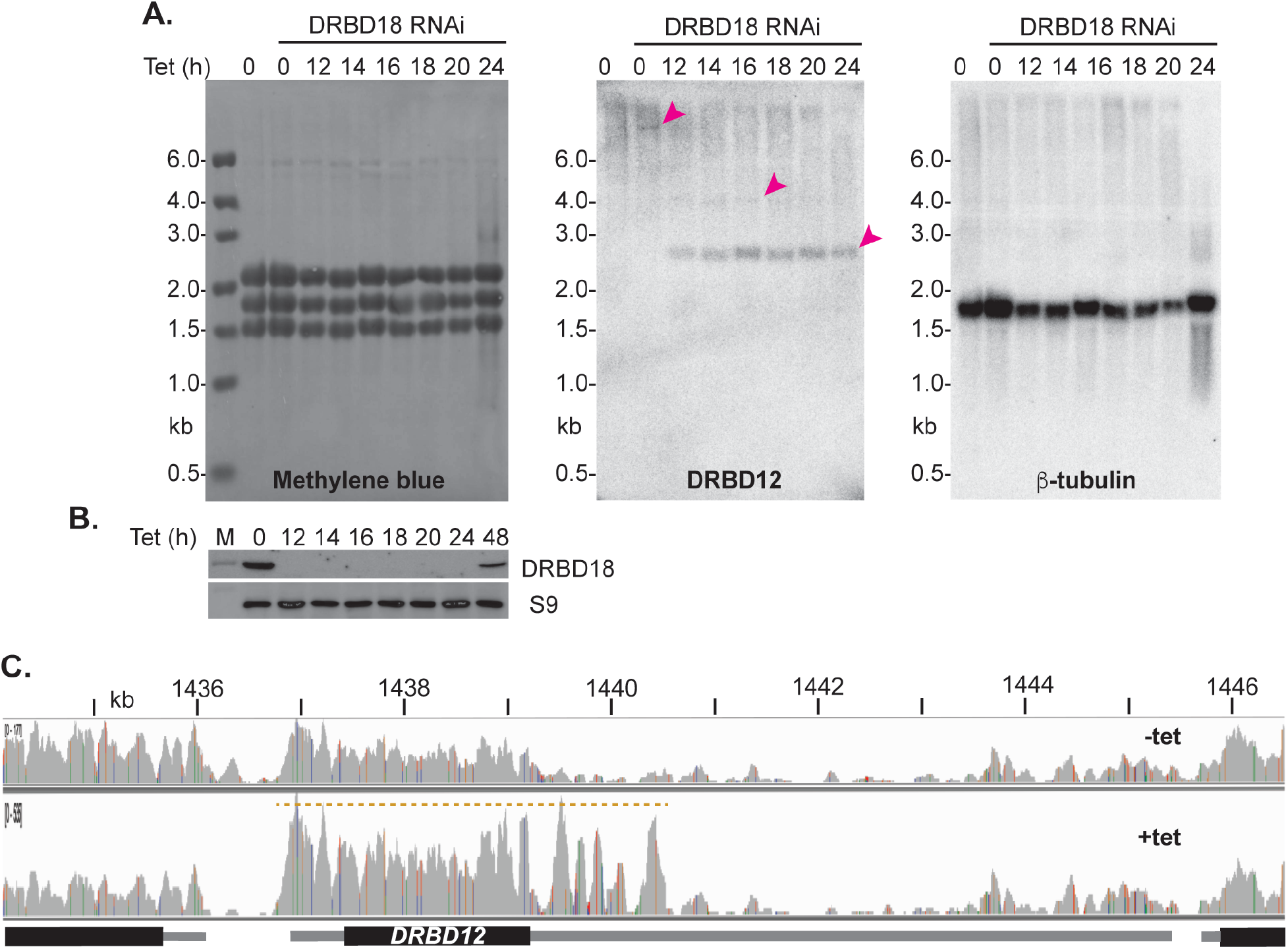
Effects of DRBD18 depletion on *DRBD12* mRNA. *DRBD18* RNAi was induced over 24h and mRNAs encoding DRBD12 and tubulin were compared, as in Figure 4. A. Northern blot results were used to examine the effects of *DRBD18* RNAi on *DRBD12* mRNA. The methylene blue-stained membrane is shown on the left, with *DRBD12* hybridisation in the centre. The membrane was then stripped and hybridised with a beta-tubulin probe as a further control. Arrows indicate the different *DRBD12* signals. B. DRBD18 protein levels after RNAi induction, corresponding to panel (A). Ribosomal protein S9 is the loading control. C. Reads mapped over the *DRBD12* region of the genome, without tetracycline or after 12h tetracycline addition: one replicate is shown but two others looked similar. Since the genomics viewer shows raw reads, the heights of the “+tet” rows have been adjusted to account for (i) different scales in the viewer and (ii) the difference in total read coverage for the two samples. This ensures that mRNAs that were not affected look approximately similar. The panel predicts a *DRBD12* mRNA of about 4 kb after 12h RNAi (orange dotted line). The low level of mapped reads over the centre of the 3’-UTR might be caused by repetitive and low-complexity sequences.

To find out how general these effects were, we further examined the transcriptomes of cells that had undergone DRBD18 depletion for 12 hours (Supplementary Table S3). The reads had been aligned to the reference TREU927 genome, and we had counted reads in both coding regions and 3’-untranslated regions. The results for the 3’-untranslated regions are not very reliable for two reasons: firstly, they contain many low-complexity sequences that may be repeated elsewhere in the genome, and more importantly, the polyadenylation site mapping in the database is unreliable. Many genes have no annotated 3’-UTRs, and the annotated ones are quite often shorter than those suggested by visual examination of the read densities.

The effects of 12h DRBD18 depletion on mRNAs in bloodstream forms were less marked than those seen after 19h induction in procyclic forms (59), and overall the two datasets correlated rather poorly (Supplementary Figure S2A). Nevertheless, there was significant overlap. Considering a set of “unique” genes (excluding paralogues (table S3, sheet 4) and *DRBD18* itself) 48 mRNAs were two-fold increased in bloodstream forms, and 243 in procyclic forms; of these, 26 were increased in both (Fisher test P-value 5.2 × 10^−26^) (Supplementary table S3 sheet 2). 89 mRNAs were 2-fold significantly (Padj <0.05) decreased in bloodstream forms and 37 in procyclic forms; of these, 12 were decreased in both (Fisher test P-value 1.3 × 10^−14^) (Supplementary table S3 sheet 3). The >1.5x decreased mRNAs in bloodstream forms included 12 that encode proteins implicated in glucose or glycerol metabolism, while the increased ones included several encoding RNA-binding proteins but also various mitochondrial proteins (Supplementary table S3 sheet 4). The changes in expression related to energy metabolism may partly just be a stress response (82) but could also be specific. There was no correlation between effects of DRBD18 depletion and differences between wild-type bloodstream and procyclic forms (not shown).

We were most interested in effects on mRNA processing. To investigate this, we looked for decreases in 3’-UTR reads (Supplementary table S3 sheet 7) that were not reflected in read counts from the coding region (Supplementary table S3 sheet 6). Overall, results for coding regions and 3’-UTRs correlated quite well (Supplementary Figure S2 B). However, for 155 mRNAs, the change in the 3’-UTR reads was at least 1.5-fold lower than the change in coding region reads (Supplementary Table S3, sheet 1). This list included some genes for which coding region counts increased significantly, and others with decreases. Most of the affected mRNAs (Supplementary Table S3, sheet 1) did not show any noticeable mRNA export defect in procyclic forms, as judged by comparing whole cell and cytoplasmic RNA (59). To verify the results for some of these mRNAs, we examined the read distributions manually. We confirmed apparent changes in processing for most (though not all) of the genes analysed: some examples are in Supplementary Figure S3. The changes affected both long and short 3’-UTRs and coding regions of various lengths. We downloaded the 200nt downstream of some of the new processing sites to look for enriched features but none were detected. Currently, therefore, we do not know why particular mRNAs are selected as DRBD18 targets.

From these results, we concluded that DRBD18 depletion had affected the processing of over 100 different mRNAs. Sometimes this resulted in increased abundance of the altered coding transcript, and more rarely, a decrease, but often there was no change.

### *DRBD18* mRNA binding does not correlate with RNAi effects

To find mRNAs that are bound to DRBD18, we purified the TAP-tagged protein, released the protein with TEV protease, and sequenced the bound RNAs. This “RIP-Seq” experiment identified 226 mRNAs that were at least 2-fold enriched (bound versus unbound) in all three purifications (Supplementary Table S6). There was no overall correlation between binding of mature transcript by DRBD18 and effects on transcript abundance (Supplementary Figure S2 C), and no correlation with mRNA length (not shown). The three values (bound/unbound) for *RBP10* mRNA were 1.6, 2.4 and 5.0; and for DRBD12 they were 2.7, 2.7 and 11.7. 226 mRNAs were reproducibly bound by DRBD18 (more than 2-fold enrichment and read counts >10 in all replicates) and of those, 36 were at least 1.5-fold changed by DRBD18 depletion - 26 increased, 10 decreased. The 3’-UTRs of these mRNAs were analysed for motifs search using the MEME suite of programmes (65). The mRNAs that were not bound by DRBD18 (< 1x-fold) and not affected were used as a background control. Two 8-nt motifs consisting of a polypyrimidine tract interrupted by a purine base (Supplementary Figure S2 D) were enriched in the bound fraction. This is interesting, since polypyrimidine tracts are needed for splicing, but this type of sequence is also found in many other 3’-UTRs. Also, the significance of the binding results is somewhat questionable. Of the 155 mRNAs that showed differences between 3’-UTR and coding region reads, only 13 were 2-fold enriched in the DRBD18 pull-down. We therefore suspect that in most cases, DRBD18 that affects mRNA processing in the nucleus - presumably by binding to processing sites - is lost either during or immediately after export of the mRNA to the cytoplasm.

### DRBD18 depletion has polar effects on polycistronic transcription units

While examining the RNAi results, we noticed that some of the down-regulated genes appeared to be in clusters on the chromosomes. Most clusters were towards the ends of transcription units, but most transcription units were not affected (Figure 9). Manual examination of the affected regions did not reveal any common features: in particular, although an mRNA with obviously altered processing was sometimes at the start of a repressed region, this was usually not obviously the case - although manual observation might not have revealed all processing defects. Conversely, genes for mRNAs with obviously altered processing were by no means always followed by a downstream region with low mRNA abundance.

**Figure 9.**
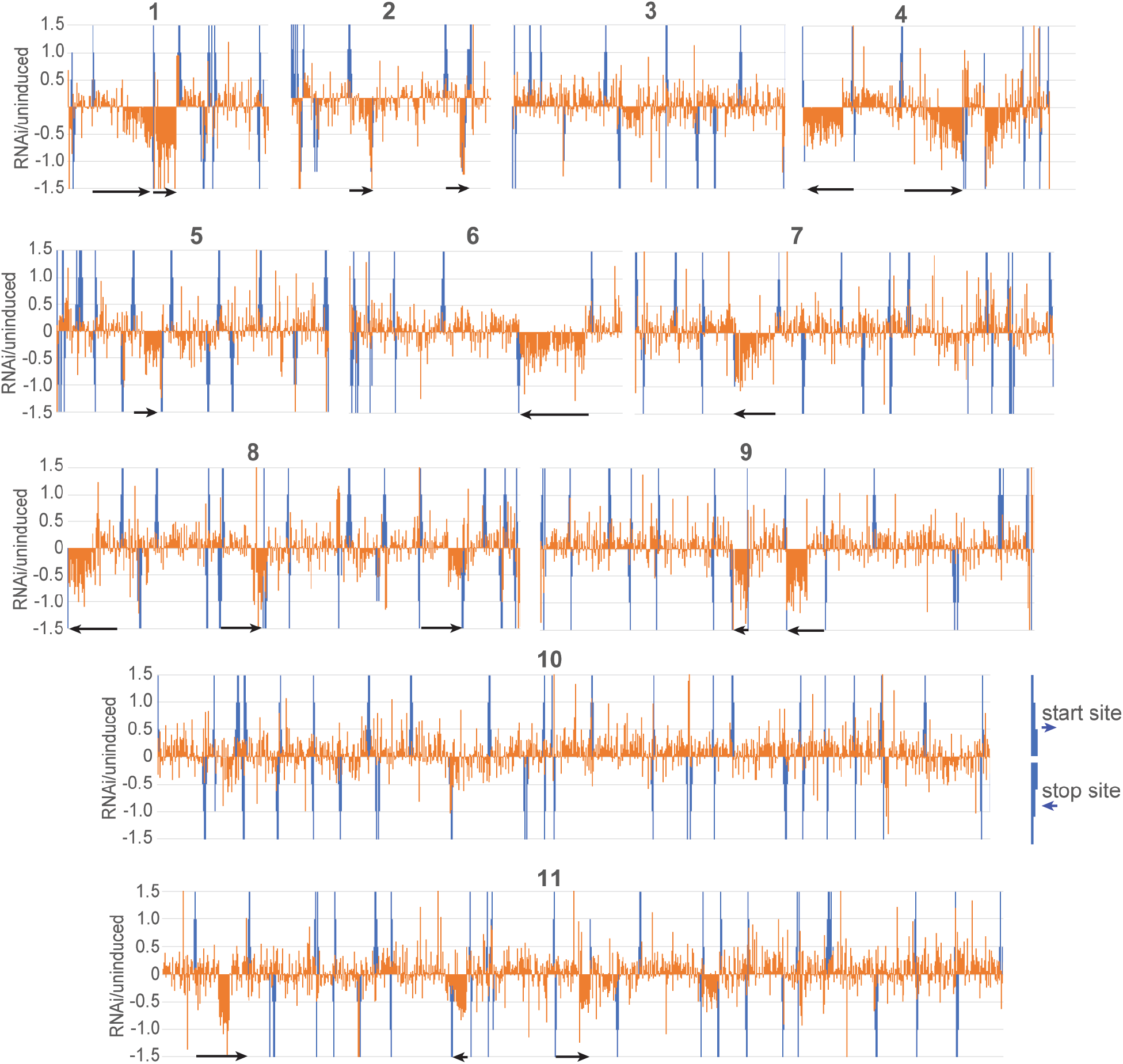
Effects of DRBD18 depletion show chromosomal clustering. The effects of DRBD18 depletion (log2 fold change) for unique genes (without paralogues) were plotted as orange bars according to the order of the genes on the chromosomes (positions not to scale). Start sites (according to histone modifications (83)) are upward blue bars and stop sites are downward bars. Neighbouring smaller bars indicate the orientation. The scale of the y-axis has been truncated so that all effects greater than 2.8-fold are shown as log_2_ (1.5). Black horizontal arrows indicate the directions of selected affected transcription units.

## Discussion

We have shown that developmental regulation of RBP10 expression is mediated by multiple different sequences in its 7.3kb 3’-UTR. Khong and Parker (84) have calculated that Opisthokont mRNAs are probably bound by at least 4-18 proteins/kb. If trypanosomes are similar, the *RBP10* 3’-UTR would be predicted to bind around 70 proteins. Thus, numerous *trans*-acting factors are likely to be acting independently to ensure appropriate expression of RBP10. In order to find proteins responsible for developmental regulation, it would be necessary to focus on individual short segments of the sequence. Our initial attempts to find effects of depleting individual proteins on RBP10 expression were therefore doomed probably to failure. We did not look for proteins that might cause *RBP10* mRNA degradation in procyclic forms, apart from DRBD18; existing results from investigations of the procyclic-specific RNA-binding proteins ZC3H20, 21 (85,86) and 22 (66) indicate that these three proteins are not involved. In particular, ZC3H20 and 21 are more likely to have stabilizing functions (85,86).

Although we did not find any proteins that are directly implicated in regulating *RBP10* mRNA stability, our results concerning DRBD18 yielded interesting insights into control of mRNA processing. Loss of DRBD18 caused retention of long *RBP10* mRNA in the nucleus, and accumulation of shorter variants which were exported to the cytoplasm. A model for a possible mechanism is shown in Figure 10 and similar effects are likely to be seen for the other mRNAs whose processing is affected by DRBD18 depletion. The *RBP10* 3’-UTR contains numerous polypyrimidine tracts of varying strengths (as judged by the number of contiguous U residues) (Supplementary text 1 and Figure 4D). We speculate that under normal conditions, DRBD18 binds to some of these (Figure 10A, (1)) and prevents binding of the *trans* spliceosome/polyadenylation complex. The interrupted pyrimidine tracts that were found to be enriched in the bound RNAs was consistent with that - although the consensus sequences found are common in most trypanosome untranslated regions.

**Figure 10.**
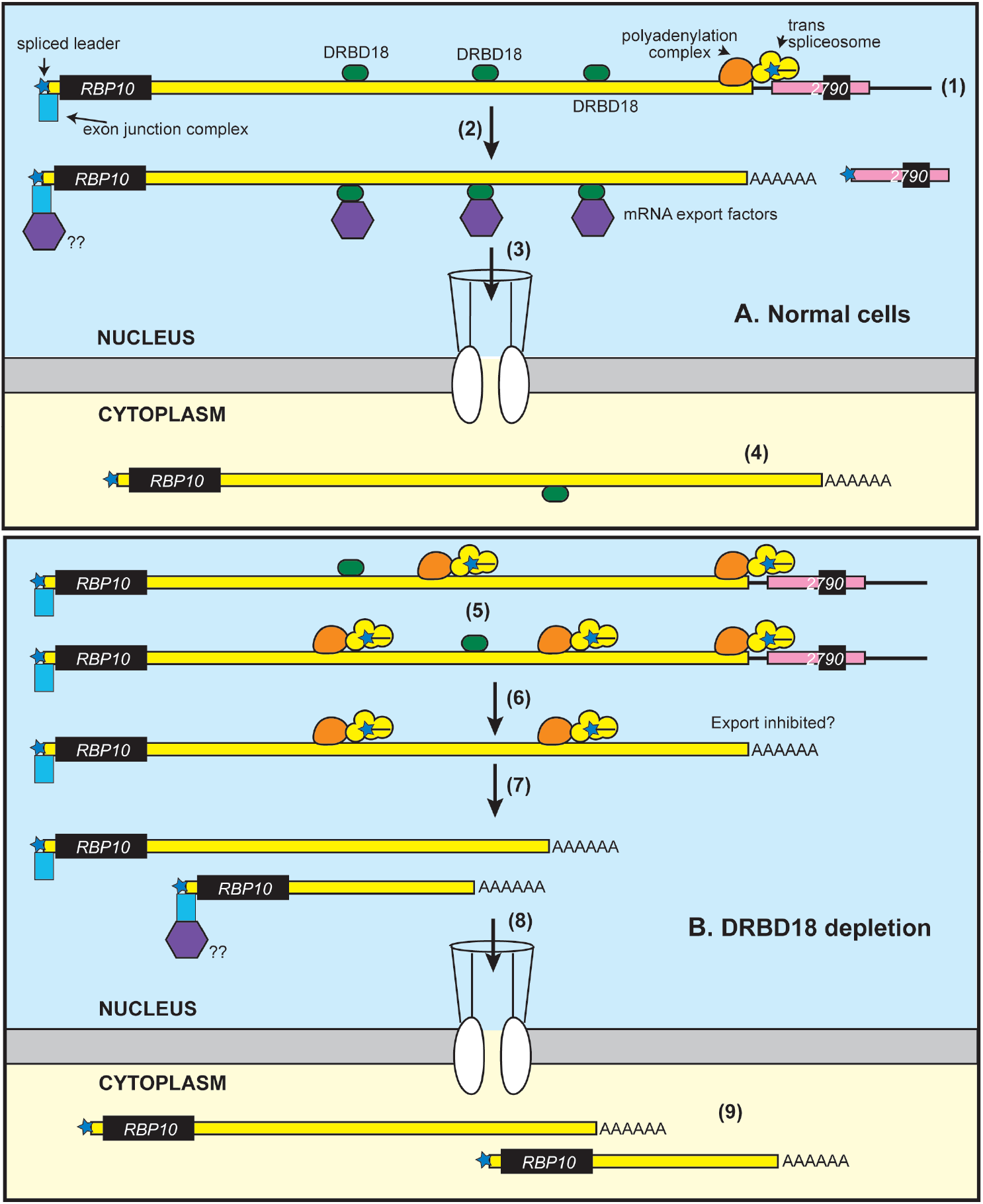
Model for the interaction between DRBD18 and *RBP10* mRNA.

We suggest that as a consequence of DRBD18 binding to the alternative sites within the 3’-UTR, processing results predominantly (though not exclusively) in the very long *RBP10* mRNA (Figure 10A, (2)). This mRNA is then exported, stimulated by interactions between DRBD18 and the export machinery (Figure 10A, (3)). A few mRNAs will also be shorter, perhaps because DRBD18 fails to bind. However, despite lacking some of the 3’-UTR, these will be developmentally regulated efficiently because even the first 2kb of the 3’-UTR is sufficient to ensure that no protein is produced in procyclic forms. Probably, the DRBD18 is lost after RBP10 mRNA export - perhaps recycling to the nucleus, since mature *RBP10* mRNA was not strongly enriched with DRBD18 (Figure 10A, (4)).

In contrast, when DRBD18 is depleted, some or all of the additional splicing signals are exposed and bound by the processing complexes (Figure 10B, (5)). The less DRBD18 that is present, the more complexes will bind. Even if the mature mRNA can be made (Figure 10B, (6)), it is retained in the nucleus. Perhaps the sequences that are concealed by DRBD18 binding bind to alternative factors that cause nuclear retention: this might include the processing machinery. mRNA export can initiate before polyadenylation has occurred (87), so precursors might be also partially outside and partially retained inside. After final processing (Figure 10B, (7)), the mRNAs can be fully exported (Figure 10B, (8)). In some cases, this might be directed by remaining DRBD18. We do not know how mature mRNAs are normally recognized for export, but since they are exported 5’-end first (87), it is possible that the exon junction complex is recognized (Figure 10). The basis for selection of mRNA precursors by DRBD18 is also unclear. In humans, the sequences bound by different RNA-binding proteins often overlap, so it has been suggested that the context - either of sequence, or of secondary structure - plays important roles (88).

Intriguingly, loss of DRBD18 caused polar decreases in the amounts of mRNA across some polycistronic transcription units (Figure 9). With the existing data, we could find no clear correlation between processing changes and downstream effects: the beginnings of these clusters did not always show an obvious processing change, and many changes in processing (including that for RBP10) did not result in downstream RNA depletion. Moreover, RNA polymerase II is likely to be at least 1 kb away before splicing occurs (89). Nevertheless, these results suggest previously unsuspected links between RNA polymerase II transcription and mRNA processing or export.

## Materials and methods

### Trypanosome culture

The experiments in this study were carried out using monomorphic *T. brucei* Lister 427 (90); except for differentiation experiments where the pleomorphic cell lines EATRO 1125 were used (91). All these cell lines constitutively express the tetracycline repressor. The bloodstream form parasites were cultured as routinely in HMI-9 medium supplemented with 10% heat inactivated foetal bovine serum at 37°C with 5% CO_2_. During proliferation, the cells were diluted to 1×10^5^ cells/ml and maintained in density between 0.2-1.5×10^6^ (92). To preserve the pleomorphic morphology between the experiments, the EATRO 1125 cells were maintained in HMI-9 medium containing 1.1% methylcellulose (93). For generation of stable cell lines, ∼1-2 × 10^7^ cells were transfected by electroporation with 10 µg of linearized plasmid at 1.5 kV on an AMAXA Nucleofector. Selection of newly transfectants was done after addition of appropriate antibiotic and serial dilution. The differentiation of bloodstream forms to procyclic forms was induced by addition of 6mM *cis*-aconitate (Sigma) to 1×10^6^ long slender trypanosomes; after 17h, the cells were transferred into the procyclic form media (∼ 8×10^5^ cells/ml) and maintained at 27°C for at least one week until they start dividing as procyclic cell lines. The stable procyclic forms were cultured as well at 27°C in MEM-Pros medium supplemented with 10% heat inactivated foetal bovine serum without CO_2_. The induction of RNAi was done using 100 ng/ml tetracycline, in the absence of antibiotics. Cells were collected after 12, 14, 16, 18, 20, 24 and 48h post-induction of the RNAi.

### Plasmid constructs

To assess the role of the *RBP10* 3’-UTR full length, a cell line expressing the chloramphenicol acetyltransferase (CAT) under the regulation of the *RBP10* 3’-UTR was generated by replacing one allele of the RBP10 coding sequence with the coding region of the CAT reporter. The *RBP10* 5’-UTR was also replaced with *ß-tubulin* 5’-UTR (Figure 1E). To map the regulatory sequences, plasmids used for stable transfection were based on pHD2164, a bicistronic vector containing a CAT and a neomycin phosphotransferase (NPT) resistance cassette genes (Figure 2A). Downstream of the CAT gene were cloned different fragments of the *RBP10* 3’-UTR in place of the *actin* 3’-UTR using *Sal* I and *Xho* I restriction sites (Figure 2A). The different fragments were obtained by PCR using genomic DNA from monomorphic bloodstream forms Lister 427 as template. Mutations on smaller fragments of the *RBP10* 3’-UTR were done using either site directed mutagenesis (NEB, Q5® Site-Directed Mutagenesis Kit Quick Protocol, E0554) or by PCR mutagenesis with Q5 DNA polymerase.

The precise details of the different constructs and their associated primers used for cloning are included in Supplementary table S7. For endogenous tagging of DRBD18, a cell line with *in-situ* TAP-DRBD18 was generated by replacing one endogenous copy of DRBD18 with a gene encoding a C-terminally TAP-tagged DRBD18. For that, a construct with neomycin resistance gene plus TAP tag cassette was flanked on the 3’-end with a fragment of *DRBD18* 3’-UTR. Upstream on the 5’-end, the C-terminal region of the *DRBD18* ORF without the stop codon was cloned in frame with the TAP tag. Prior to transfection in monomorphic bloodstream forms Lister 427, the plasmid (pHD3200) was cut with *Apa* I and *Xba* I to allow homologous recombination. The tetracycline inducible construct for DRBD18 RNAi was done using a stem-loop vector targeting the coding region of DRBD18. The region used for the stem loop vector was checked against off-targets using the RNAi target selection tool for trypanosome genomes (https://dag.compbio.dundee.ac.uk/RNAit/) as described in (94).

### RNA analysis and Northern Blotting

Total RNA was isolated from approximately 1×10^8^ bloodstream-form trypanosome cells or 5×10^7^ procyclic-form cells growing in logarithmic phase using either peqGold Trifast (PeqLab) or RNAzol RT following the manufacturer’s instructions. To detect the mRNA by Northern blot, 5 or 10 µg of the purified RNA was resolved on formaldehyde agarose gels and transferred onto nylon membranes (GE Healthcare) by capillary blotting and fixed by UV-crosslinking. The membranes were pre-hybridized in 5x SSC, 0,5 % SDS with 200 mg/ml of salmon sperm DNA (200 mg/ml) and 1x Denhardt’s solution, for an hour at 65°C. The probes were generated by PCR of the coding sequences of the targeted mRNAs, followed by incorporation of radiolabelled [α^32^P]-dCTP and purification using the QIAGEN nucleotide removal kit according to the manufacturer’s instructions. The purified probes were then added to the prehybridization solution and the membranes were hybridized with the respective probes at 65°C for overnight (while rotating). After rinsing the membranes in 2x SSC buffer/0.5% SDS twice for 15 minutes, the probes were washed out once with 1x SSC buffer/0.5% SDS at 65°C for 15 minutes and twice in 0.1x SSC buffer/0.5% SDS at 65°C each for 10 minutes. The blots were then exposed onto autoradiography films for 24-48 hours and the signals were detected with the phosphorimager (Fuji, FLA-7000, GE Healthcare). The signal intensities of the images were measured using ImageJ.

Total RNA used for sequencing analysis (RNA-Seq) was prepared from cells collected 12 hours after the induction of DRBD18 RNAi in monomorphic bloodstream forms cells (Lister 427). Induction of RNAi was done using tetracycline at a concentration of 100 ng/ml. The cells without induction of DRBD18 RNAi were used as controls. RNA was extracted using the phenol-chloroform separation method as described previously. The integrity of the total RNA was checked on a denaturing agarose gel. Afterwards, the ribosomal RNAs (rRNAs) was depleted from the total RNA samples using a cocktail of oligonucleotides complementary to rRNAs as previously described (17,95). The purified samples were then submitted to the Core Facility for Deep Sequencing.

### RNA half-life assay

mRNA transcription and *trans*-splicing were simultaneously inhibited by addition to the growth culture medium of 10 µg/ml Actinomycin D and 2 µg/ml Sinefungin. The cells were collected at the indicated different time points and RNA was isolated by Trizol extraction (68). The mRNA levels were assessed by Northern blotting using [α^32^P]-radiolabelled probes, and quantitated by phosphorimaging. The signal intensities of the images were measured using ImageJ.

### CAT Assay

To perform the CAT assay experiment, approximately 2×10^7^ cells expressing the CAT reporter gene were harvested at 2300 rpm for 8 minutes and washed three times with 1X cold PBS. The pellet was re-suspended in 200 μl of CAT buffer (100mM Tris-HCl pH 7.8) and lysed by freeze-thawing three times using liquid nitrogen and a 37°C heating block. The supernatants were then collected by centrifugation at 15,000×g for 5 min and kept in ice. The protein concentrations were determined by Bradford assay (BioRad) according to the manufacturer’s protocol. For each setup, 0.5 μg of protein in 50 μl of CAT buffer, 10 μl of radioactive butyryl CoA (^14^C), 2 μl of chloramphenicol (stock: 40 mg/ml), 200 μl of CAT buffer and 4 ml of scintillation cocktail were mixed in a Wheaton scintillation tube HDPE (neoLab #9-0149) and the incorporation of radioactive acetyl group on chloramphenicol was measured using a Beckman LS 6000IC scintillation counter.

### RNA immunoprecipitation

A cell line expressing *in-situ* C-terminally TAP tagged DRBD18 was used for the RNA immunoprecipitation. The TAP-tag consists of the protein A and calmodulin binding protein domains separated by a Tobacco Etch Virus (TEV) protease cleavage site. Approximately 1×10^9^ cells expressing *in-situ* C-TAP DRBD18 with a concentration of 11×10^6^ cells/ml were pelleted by centrifugation at 3000 rpm for 13 minutes at 4°C. The pellet was washed twice in cold 1x PBS and collected by centrifugation at 2300 rpm for 8 minutes at 4°C and then snap frozen in liquid nitrogen. The cell pellet was lysed in 1 ml of the lysis buffer (20 mM Tris pH 7.5, 5 mM MgCL_2_, 0.1% IGEPAL, 1 mM DTT, 100 U RNAsin, 10 μg/ml leupeptin, 10 μg/ml Aprotinin) by passing 20 times through a 21G x ½ needle using a 1 ml syringe and 20 times through a 27G x ¾ needle using a 1 ml syringe. The lysate was cleared by centrifugation at 15,000 g for 15 minutes at 4°C. Afterwards, the supernatant was transferred to a new Eppendorf tube and the salt concentration was adjusted to 150 mM KCl. 10% of lysate was collected as the input fraction for RNA extraction and 2% was used for western blotting. 40 μl of magnetic beads (Dynabeads M-280 Tosylactivated, Invitrogen) coupled with Rabbit Gamma globulin antibodies (Jackson Immuno Research Laboratories) were washed three times with 500μl of IP buffer (20 mM Tris pH 7.5, 5 mM MgCL_2_, 150 mM KCl, 0.1% IGEPAL, 1 mM DTT, 100 U RNAsin, 10 μg/ml leupeptin, 10 μg/ml Aprotinin) then incubated with the cell lysate at 4°C for 3 hours with gentle rocking. The unbound fraction was collected for RNA extraction (98%) and Western blotting (2%). Subsequently, the beads were washed thrice with IP buffer. 2% of each wash fraction was collected for protein detection. The TAP-tag was cleaved by incubating the beads with 500 μl of IP buffer containing 100 units of TEV protease (5 μl) with gentle rotation at 16°C for 2 hours. The eluate was then collected afterward by magnetic separation for RNA isolation and protein detection. RNA was isolated from the input, the unbound and the eluate fraction using the peqGOLD Trifast FL (Peqlab, GMBH) according the manufacturer’s instructions. To assess the quality of the purified RNA, aliquots of the input, unbound and eluate fractions were resolved on formaldehyde agarose gels to check the integrity of the ribosomal RNAs. Total RNA from the unbound and the eluate fraction were depleted of ribosomal RNA (rRNA) using RNAse H (NEB, M0297S) and a cocktail of 131 DNA oligos (50 bases) complementary to the trypanosome rRNAs (95). Following rRNA depletion, the samples were subjected to DNAse I treatment in order to remove any trace of oligonucleotides using the Turbo DNAse kit (Invitrogen, ThermoScientific). The RNA samples were afterward purified using the RNA Clean & Concentrator -5 kit (ZYMO RESEARCH) following the manufacturer’s instructions. The recovered purified RNA from both bound and unbound samples was then analysed by RNA-Seq.

### High throughput RNA sequencing and bioinformatic analysis

RNA Sequencing was performed at the Cell Networks Deep Sequencing Core Facility of the University of Heidelberg. The library preparation was done using the NEBNext Ultra II Directional RNA Library Prep Kit for Illumina (NEB, E7760S). The libraries were multiplexed (six samples per lane) and sequenced with a NextSeq 550 system, generating single-end sequencing reads of about 75 bp. All analysis was done using a custom pipeline (96,97). Before analysis, the quality of the raw sequencing data was checked using FastQC (http://www.bioinformatics.babraham.ac.uk/projects/fastqc). Cutadapt (98) was used to remove sequencing primers, then the sequencing data were aligned to *T. brucei* 927 reference genome using Bowtie (99) allowing 1 alignments per read, then sorted and indexed using SAMtools (100). For RNAi, the reads that mapped to the open reading frames, 3’-UTRs and non-coding RNAs in the TREU 927 genome were counted.

In order to look for enrichment of particular functional characteristics, we used a list of unique genes modified from (12); this corrects for repeated genes and multigene families. To generate reads per million reads for the unique gene set, we first extracted the list, then multiplied the reads by the gene copy number (obtained in a separate analysis, see e.g. (101)). For the RIP-Seq, the reads per millions were counted and the ratios eluate versus flow-through were calculated. An mRNA was considered as “bound mRNA” if the ratios from all the three pulldowns were higher than 1.5. The 3’-UTR motif enrichment search was done using MEME in the relative enrichment mode (64). Annotated 3’-UTRs were downloaded from TritrypDB. For some other 3’-UTRs, the manual annotation was performed using the RNA-Seq reads (e.g. (15) and poly(A) site data in TritrypDB (102). Analysis of differentially expressed genes after DRBD18 RNAi was done in R using DESeqUI (96), a customized version of DESeq2 (103) adapted for trypanosome transcriptomes. Statistical analyses were done using R and Microsoft excel. Raw data are available at Annotare as E-MTAB-9783 and E-MTAB-10735.

### Western blotting

Protein samples were collected from approximately 5×10^6^ cells growing at logarithmic phase. Samples were run according to standard protein separation procedures using SDS-PAGE. The primary antibodies used in this study were: rabbit α-DRBD18 (1:2500, (58)), rat α-RBP10 (1:2000, (28)); rat α-ribosomal protein S9 (unpublished) and rabbit anti-XRND (1:2500 (68)). We used horseradish peroxidase coupled secondary antibodies (α-rat, 1:2000 and α-rabbit, 1:2500). The blots were developed using an enhanced chemiluminescence kit (Amersham) according to the manufacturer’s instructions. The signal intensities of the images were quantified using ImageJ.

### Protein-protein interactions

A cell line in which one allele of DRBD18 bear a sequence encoding a C-terminal TAP tag was used to study the interactome of DRBD18. TAP-DRBD18 was purified using IgG magnetic beads and the bound protein was eluted with TEV protease, which was then removed using HisPur Ni-NTA magnetic beads (Thermo Scientific) according to the manufacturer’s instructions (104). The proteins from three independent purifications were subjected to denaturing SDS-PAGE and analysed by mass spectrometry. A cell line inducibly expressing the GFP with a TAP tag at the N-terminus was used as a control. The expression of the GFP-TAP was induced with 100 ng/ml tetracycline for 24 hours and the purification of the protein was done as for C-TAP DRBD18. After purification, the proteins were subjected to denaturing SDS-PAGE, visualized by Coomassie blue staining and analysed by mass spectrometry (76). Data were quantitatively analysed using Perseus (72).

### Subcellular fractionation

Nuclear-cytosolic fractionation was performed as described in (105). Briefly, 3×10^8^ Lister 427 bloodstream form wild-type cells as well as those expressing the RNAi stem loop vector targeting DRBD18 were harvested by centrifugation at 3000 rpm for 13 minutes at 4°C. The cell pellet was washed twice with cold 1x PBS and collected by centrifugation at 2300 rpm for 8 minutes at 4°C. Afterwards, the pellet was resuspended in hypotonic buffer (10 mM HEPES pH 7.9, 1.5 mM MgCl2, 10 mM KCl, 0.5 mM dithiothreitol, 5 μg/ml leupeptin and 100 U RNasin) and lysed in presence of 0,1% IGEPAL (Nonidet P-40) by passing 20 times through a 21G x ½ needle using a 1 ml syringe and 20 times through a 27G x ¾ needle using a 1 ml syringe. The cells were allowed to rest on ice for 20 minutes. Then, the nuclei fraction was pelleted by centrifugation at 15,000 g for 15 minutes at 4°C. The supernatant containing the cytoplasmic fraction was transferred to a new tube. 20 % of the cytoplasmic fraction was used for western blotting and the rest was used for RNA extraction. The pellet containing the nuclei fraction was resuspended in TBS containing 0.1 % SDS; 20 % of this fraction was used for western blotting while the rest was used for RNA extraction (result not shown).

### Affymetrix smRNA FISH

The Affymetrix single molecule mRNA FISH was carried out as described in (81). A total of 200 ml bloodstream-form trypanosomes at ∼ 5-8 ×10^5^ cells/ml were harvested by centrifugation (8 minutes, 1400g), resuspended in 1 ml 1x PBS and pelleted again by centrifugation (5 minutes, 1400g). The cell pellet was resuspended in 1 ml 1x PBS, followed by fixation with 1 ml of 8% formaldehyde (in PBS). The mixture was incubated at room temperature for 10 minutes with an orbital mixer. A total of 13 ml 1x PBS were added and the cells were harvested by centrifugation (5 minutes, 1400 g). The pellet of fixed cells was resuspended in 1 ml 1x PBS and spread on glass microscopy slides (previously incubated at 180°C for 2h for RNAse removal) within circles of hydrophobic barriers (PAP pen, Sigma). The cells were allowed to settle at room temperature for 20 minutes. The slides were then washed twice in 1x PBS. The protease solution was diluted 1:1000 in 1x PBS and briefly vortexed to allow complete dissolution. Permeabilization of the fixed cells was done with addition of 50 μl of detergent solution QC in each circle on the slides. This was followed by a 2-step washing in 1x PBS. 100 μl of the protease solution was added to each circle and incubated exactly for 10 minutes at 25°C. The slides were then washed twice in 1x PBS and used for Affymetrix FISH experiments as described in the manual of the QuantiGene ViewRNA ISH Cell Assay (Affymetrix), protocol for glass slide format. The only modification from the kit protocol is that the protease digestion was done at 25°C rather than the normal room temperature and we used a self-made washing buffer (0.1x SSC buffer, 0,1% SDS) instead of the washing buffer from the kit. All Affymetrix probe sets used in this work are described in supplementary Table S7 (sheet 3). For visualization, the labelled cells were mounted with 4′,6-diamino-2-phenylindole dihydrochloride (DAPI) solution, diluted 1:1000 in 1x PBS. Images were taken with a fluorescent inverted wide-field microscope Leica DMI6000B (Leica Microsystems GmbH, Wetzlar, Germany) equipped with 100x oil immersion (NA 1.4) and a Leica DFC365 camera (6.45 m/pixel). Deconvolution was done using Huygens Essential software (SVI, Hilversum, The Netherlands) and images are presented as Z-stack projection (sum slices). The image analysis was carried out using the available tools in Image J software and 50 cells for each slide were selected for quantifying the number of mRNAs present in the cytosol and in the nucleus.

## Data availability

Raw RNASeq data are available at Annotare as E-MTAB-9783 and E-MTAB-10735. The mass spectrometry proteomics data have been deposited to the ProteomeXchange Consortium via the PRIDE partner repository (106) with the dataset identifier PXD027792.

## Author contributions

Tania Bishola was responsible for nearly all of the experimental work, provided figures and tables, wrote the first draft of the paper and was involved in subsequent editing. Bin Liu supervised in the initial phase of the study. Christine Clayton devised and supervised the project, edited the paper and provided funding.

## Acknowledgments

We are very grateful to Dr. Susanne Kramer, who supervised and provided facilities for the mRNA FISH experiment. We thank Dr. Monica Terrao for some initial analysis of the *RBP10* 3’-UTR. Six fragments of the *RBP10* 3’-UTR were cloned by Juyeop Kim, a master student, during his 6-week lab rotation. We are thankful to Prof. Dr. Laurie Read for antibody donation and for extensive and friendly discussions, including communicating unpublished results. We thank Bernardo Gabiatti and Paula Andrea Castañeda Londoño (Biozentrum, Universität Würzburg) for discussion and assistance with the mRNA FISH experiment; Laura Armbruster (COS, University of Heidelberg) for help in mass spectrometry data analysis. All RNA-Seq libraries and RNA sequencing were done by David Ibberson at the Bioquant in the University of Heidelberg. Mass spectrometry was done in the ZMBH Core Facility for Mass Spectrometry by Thomas Ruppert and Sabine Merker. We thank Claudia Helbig and Ute Leibfried for technical assistance, for preparing media and buffers. We are indebted to Prof. Dr. Nina Papavasiliou (DKFZ, University of Heidelberg) and Prof. Dr. Luise Krauth-Siegel (BZH, University of Heidelberg) for allowing us to share their laboratories including equipment and reagents after the flood in ZMBH.

## SUPPLEMENTS

**Supplementary Table S1. Sizes of reporter *CAT* reporter RNAs**

Expected and observed sizes of all reporter mRNAs. The expected sizes are predicted from the sequences, with 100 nt added to allow for a 39nt spliced leader and a 61nt poly(A) tail; known poly(A) lengths can be up to about 200nt (56,107). Some mRNAs were shorter than expected, suggesting that a poly(A) site within the 3’-UTR sequences was used. Two reporters (F2.2.1 and F2.2.3) gave two different mRNA variants, one with nearly the expected size and another that was longer or shorter.

**Supplementary Table S2. List of proteins targeted by RNAi.**

**Supplementary Table S3: Effect of *DRBD18* RNAi on the transcriptome**

**Supplementary Table S4: Mass spectrometry results**

**Supplementary Table S5: Detailed FISH results**

**Supplementary Figure S1: Effect of *DRBD18* RNAi on RBP10 protein and RNA expression in EATRO1125 bloodstream and procyclic forms; cell fractionation**.

**A**. Cumulative growth curves of EATRO1125 bloodstream forms with and without induction of *DRBD18* RNAi.

**B**. DRBD18 and RBP10 protein levels after *DRBD18* RNAi induction in bloodstream forms. Wild-type cells procyclic forms were used as control.

**C**. DRBD18 and RBP10 protein levels after *DRBD18* RNAi induction in procyclic forms. (a) detection of DRBD18; (b) detection of RBP10 on the same blot, *cross-reacting band; (c) a different experiment with longer exposure. The orange arrow points to the very faint RBP10 band seen after 2 days of DRBD18 depletion. Wild-type (WT, no RNAi) controls are P: procyclic forms; B: bloodstream forms.

**D, E**. Lister 427 bloodstream-form trypanosomes were separated into nuclear and cytoplasmic fractions at various times after RNAi induction. D and E are different blots from the same experiment. The distributions of DRBD18 (panel D), RBP10 (cytoplasmic, panel D with the DRBD18 signal still visible) XRND (a soluble nuclear protein, panel E) and ribosomal protein S9 (nucleus and cytoplasm) were examined.

**F**. Effect of RNAi on *RBP10* mRNA in EATRO1125 bloodstream forms. The methylene-blue-stained blot is on the left and the signal from *RBP10* mRNA is on the right. The corresponding Western blot for DRBD18 is shown in panel G.

**Supplementary Figure S2. DRBD18 depletion and binding analysis**

**A**. The effect of DRBD18 RNAi in procyclic forms (y-axis) was plotted against the effect of DRBD18 RNAi in bloodstream forms (x-axis)

**B**. Correlation between the effect of DRBD18 depletion in bloodstream forms on coding sequences (y-axis) vs 3’-UTRs (x-axis)

**C**. Lack of correlation between DRBD18 depletion in bloodstream forms (y-axis) and DRBD18 binding (x-axis).

**D**. Motifs enriched in DRBD18 bound mRNAs relative to size-matched controls not affected by DRBD18 RNAi and showing less than 1-fold enrichment in the bound fraction.

**Supplementary Figure S3. Effects of DRBD18 depletion on mRNA processing**

DRBD18 RNAi was induced for 12h and the RNA was sequenced. The figure shows typical results for one of the three replicates (replicate 1), displayed on the Integrated Genomics Viewer (69,70). In the maps coding regions are black and non-coding regions are grey, with an arrow at the 3’-end of each mRNA. Transcription direction is from left to right. Since the genomics viewer shows raw reads, the heights of the “+tet” rows have been adjusted to account for (i) different scales in the viewer and (ii) the difference in total read coverage for the two samples. This ensures that mRNAs that were not affected look approximately similar. “Lost” 3’-UTR regions are indicated with cyan dotted lines.

A. Tb927.10.14390. B. Tb927.8.5240; C. Tb927.4.2500; D. Tb927.11.16780; E. Tb927.11.2500.

