## Supplementary figures and images for "The RNA-binding protein DRBD18 regulates processing and export of the mRNA encoding *Trypanosoma brucei* RNA-binding protein 10"

### Supplementary Figure S1

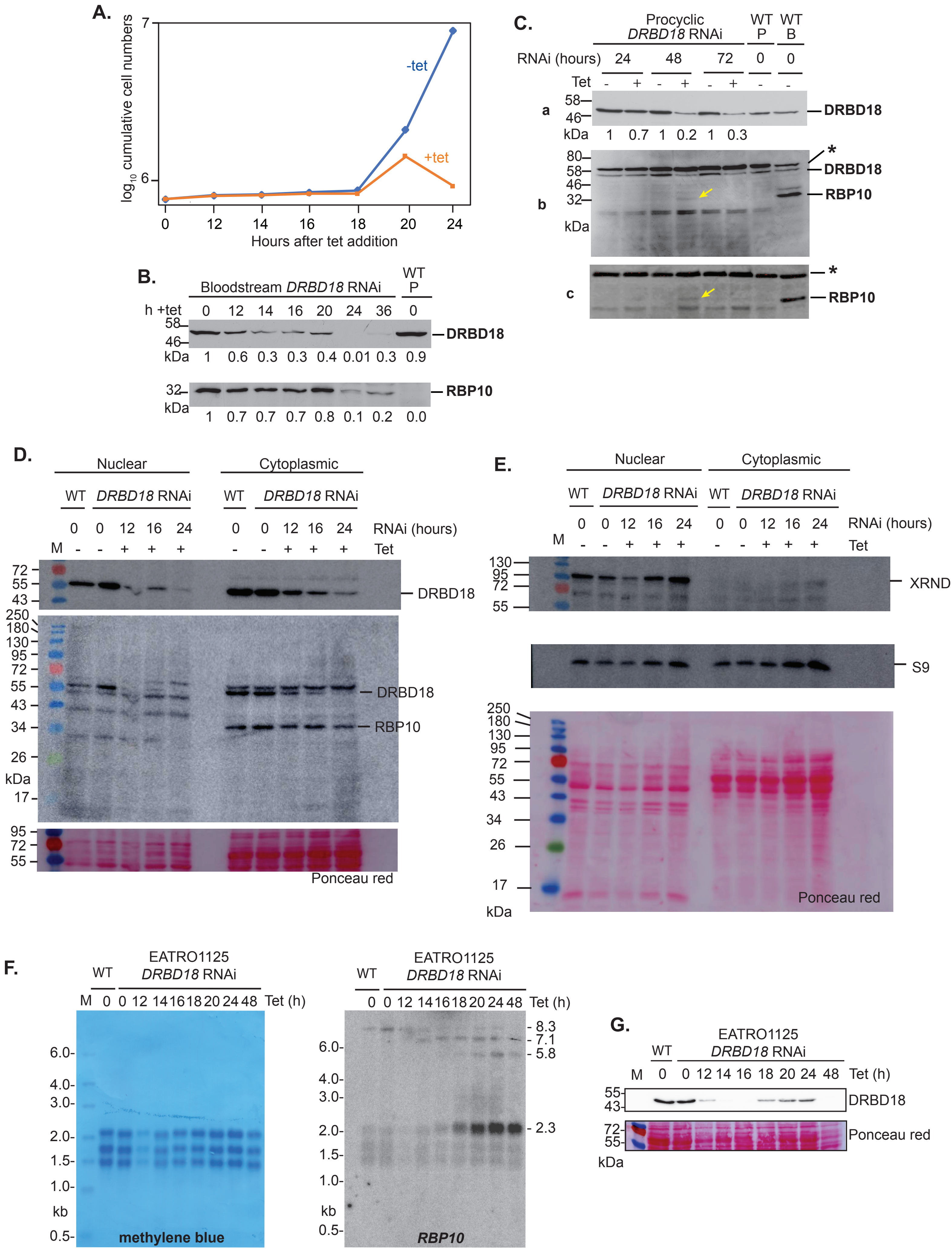

### Supplementary Figure S2

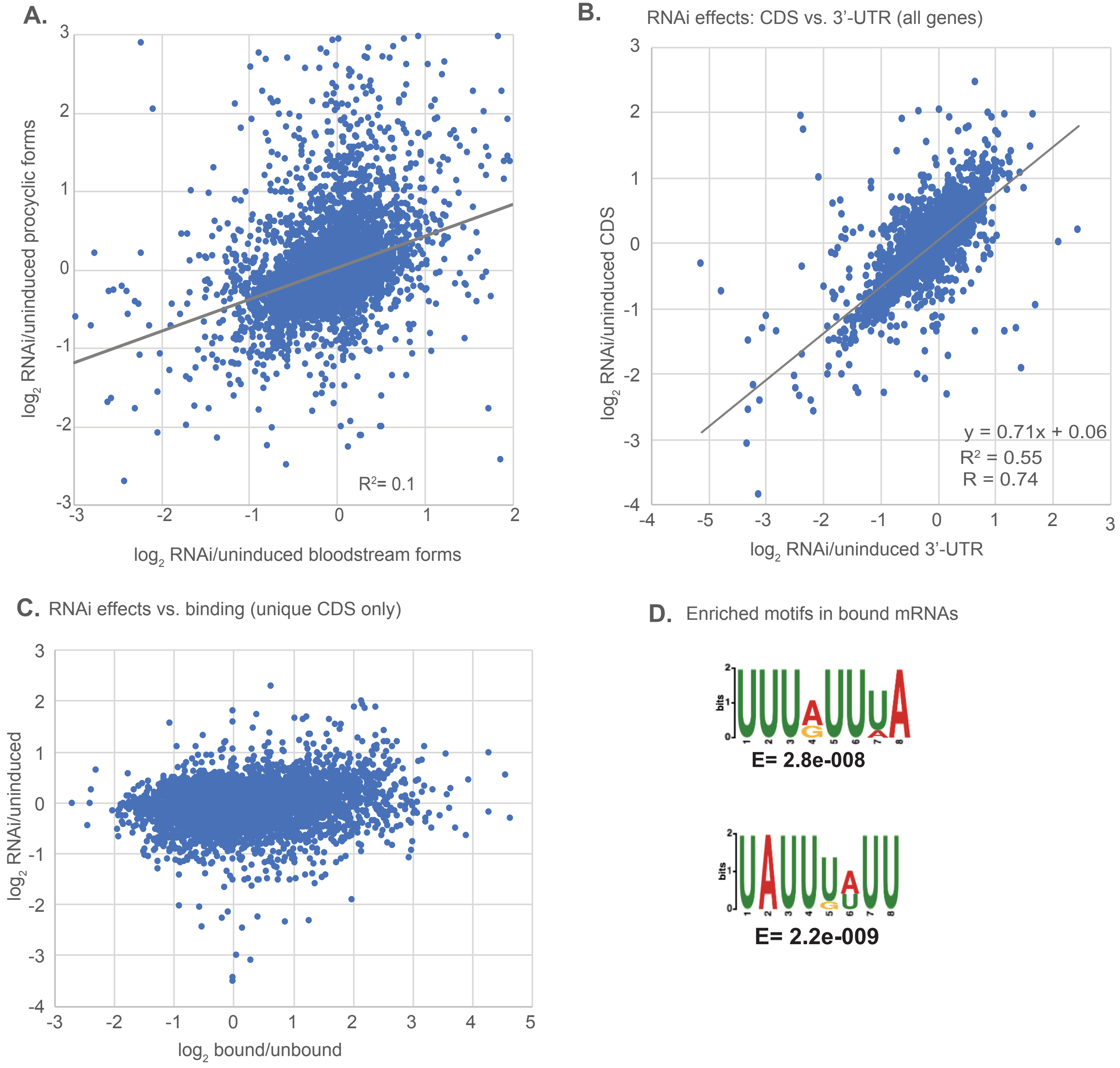

### Supplementary Figure S3

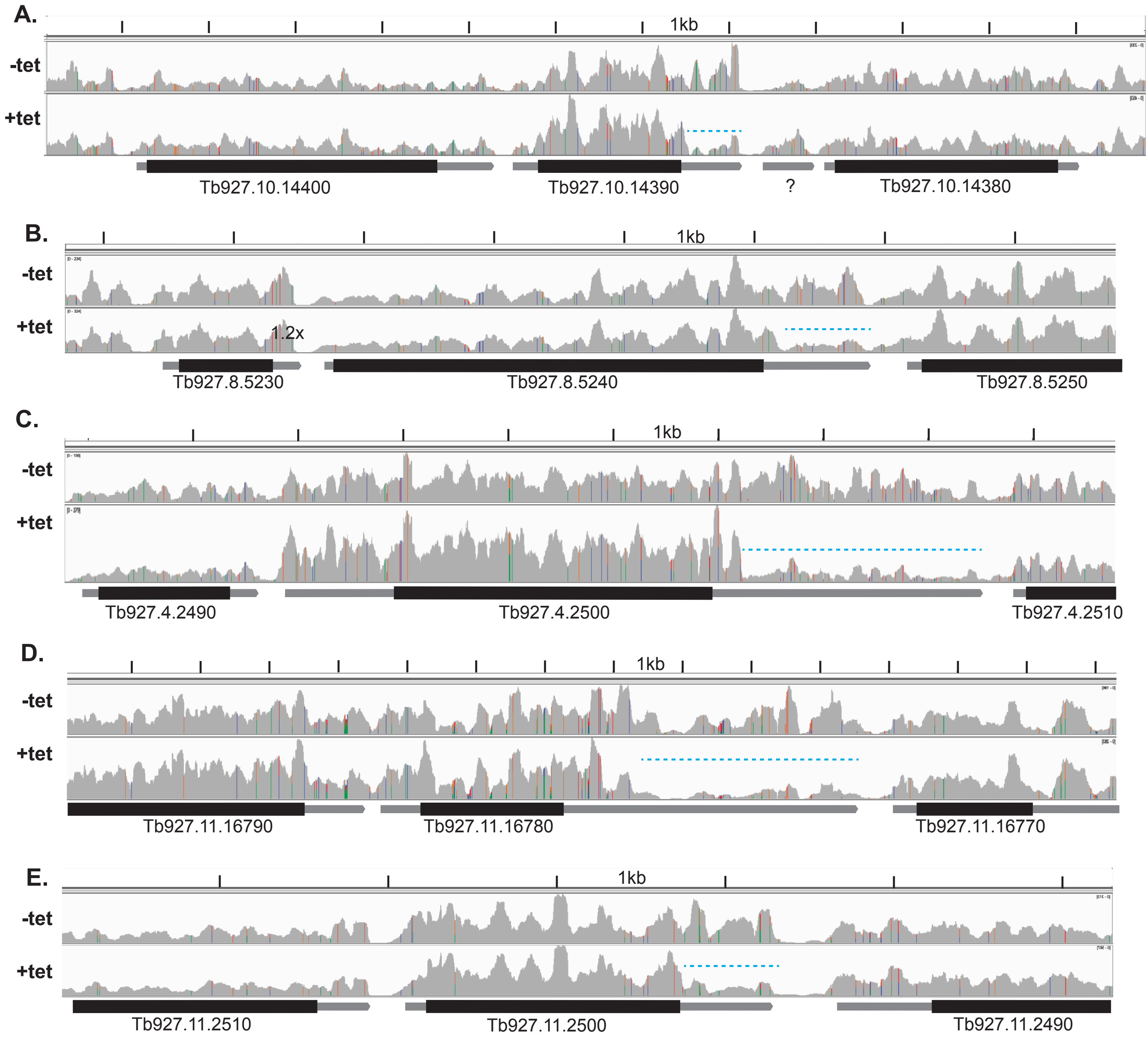
