## Supplementary Text S1 for "The RNA-binding protein DRBD18 regulates processing and export of the mRNA encoding *Trypanosoma brucei* RNA-binding protein 10"

**>Full sequence with UUUAUUUA and (U)_9_ underlined.**

**UGGCACAGAGGGUAACGAAGUAGGAAUUUUUGCCGCCAGCUGAGGUCGUUUACCUUGGGUUGGCUGUCAUGGAGAUAGGGAAGAGAGAAGCAACAUCGCGUGCAAGGAAAAACGAACAGAAGAAUCGAUCCCUCCCCCCUCCAUGAGUCCUUCUUCGUUUAUAAUUUCGGCUGUUGUUUUUGCUGUCACCUUUUUUUUUCUUCCUUCUCGUUCGCCACACCCUCUCCUCAACCUCCUUUACCUCCAUGGAUCUCUCGGCAGCAGUCCCCCUCCCCUCAUUCCCCUCAUGUUUUGGUGGUCGCCUUAUUGGCACCUCUUUUCGUCAUUUUUUCCCCCCACUUGCUGCAGUUCAAGGCGUAUUUGUGGGGAGAGUAAAGAACAUAAGUGAAAUCUGAAAACAAUAGAAAAGAGAGAUGAAUUAAUUAAUGAAUGAAUGGGAGAAAUAAGGAGGAGAGGUGGUGGAGAGCUGGGGAACUGUAAUGAAGAAGAAAAAAAAGAAGAACGAGGGAAAGGUAGAUUCAGUGGGGGAGACAAAACAGUGAAGAAGAAUCUGGCGGAAAGCAACAAUAAGUGUGAAAAUGAGAACGAAGGUGUGAAUGAUUCUUCAGUUUAAGAAACCACACGUGAAGAAGGACAGAAUAUAUAAAUAAAUAUAUCUAUUUAUAUGUUGAUGUCUCAGAAGAAAAAAAGCAAAGGAGGGGAGAUAGAAGAGGUUUAAAGGGAGGAAGGAAGACAGGCCCUGAAACCAAUAAAACAAAAUAAAUAAAUAAAGAAACGCGAAAUCAAAACUGAAAAUGCGAAAAAAAAAACGAAGAAGAAGAAGGGUGCAAUGAAGGAAUGAACUCCACAGAGAGAAUACCCGUUCAAACAUUCUUUUCUUCGUUUUUCCCCUCCUCCUCUCCCCGACGCAACCGCCCCCCUUUUUUUUCUAUAAUUGGCUCUUCUGACCCACACCCCCUCACUUUCCUUCUCAAAGAACGGCAUCUUGUUUUCCCACCCUUUAUUAUUGUUAAUUUCUUUUUUCGCUUUACUAUUAUUAUUAUUAUUUACCACCCCAUCACCGUUACUUUUAGUUGUUUUUUUUUUUUUUAUGCCUCUCUCCUUUCAUUUUUUUGCGCUUUUGCUCCUCCUUAUUGGUUAUGUAGAGCGAUUUAUAUAAAUCUAAUAUAUAUAUAUAUAUAUAUAUUUGUAUUCAUAUAUUUUAUGUCUUAUAUACAUAACUACUUUCGCGCAGAAAAGGGAAAGAGGAAAGGAGAAGAAGGGGGAAAAUAGAAAGCCCAUGCACAUAUAUAUAUAUAUAUAUUUAUUUAUUUGUUUAUUUAUUUAUAUAUAUAUAUAUAUGUAUUCUUUGUACCUCCCUUUCGUUUUUACUUUGUUUUUUGUUUUACCUUUUUUUUGUUUCUACUUUGUUUCUUGUUUUGUUUACUGUUAGUCACUCCUUACUAUUUUUACCGCCUCUAUUAUUAUCACUAUUAUUAUUAUUAUUAUUACUAUUAUUAUUCCUAUUAUUAUUAUUAUUAUUCCUAUUAUUAUUAUUAUUAUUAUUCCUAUUAUUAUUAUUAUUCCUAUUACUACUGCUAUUAUUAUUGUUAUUUCUGGGGCUACUGCAGUGCUUUCCCUUUCUUUUUCUUUGGGUGUUCGCGUGGUUGAGAUUGUGAUGUGACUUCUGUUUGCUGACGAUUGUUUGUUUCUAUUUGAUUGUGCCGGUGUCUGUUUUCCCCCACCCCCGUGAUGUCACGAAGAAAAACAAAAAUUAAAUUGAAAAGUUUUUACUUCUCCUCGUCCAAAGCAAUUGCUCUCCUCCUUCCUUUUCUCCACACGCGCGCACGUACGUAGUGAACUAAUCAAAGAGUGAAGAAAUAAAAAUAAAAACACGCGUCGUGUUGUGGAUCCCUUUUUCGGAUCCAAUUUGCGCUAUUGCUUUUUUUUGUUAGUAUUAUUGUUUUGUGCUUUUUUUUUUCUUUUUCUGUGUGUGUAAUAUGAUUUUGUCGCUUUUCAUUCAGCGGAAGCAGACAGUAAAGUUAUGGGCUCAUAUGCUUGCAUGUGCAUUCCAUCCGAUAUUGCGCGGGAGUUGUUGUUUUGUUUUGUUUUUCACACCUUCGUCCCUUUUUUUUUUCUUUUUGGUUUUUAUUGUUUGUAUAUAUUUUCAUAUUUAUGUACAUAUAUUAGGAGCAAAUGCAUGCGUGUUUGUGCUUUAACGCCCAUGUGUGCGCCCCUUGGCUUGCUUUCCCACAAAUACCCUUUCGAAACGCCUUACGGAAGCGGGGACAAUAUAAUUAAGAAGAAAGGGAGAUGGGGGGGAAAAAAAAAAGAGAAAGAGAAAGAGAAACACCGGCUCGUAGACAUGAGGAGAGGAAUCAAAAAAAAGACAAUGAAAUGAAAUAAAGUUAAGUGAAGUGAAUUGAAACGAGGUAAGAAUUUAAGAAGUUAAGAAAGCUGGUGACCCGCCGUCGCUUUUCUACUUCAUUCCACUCUUUUUUUUUCUUCUUUUCUUUUUUUUCCCAUCCUUUUCCUUUUUUUUUGCUAGUUUUUGAUCCGCUUUGCCUUUGCAUCUUAUCUUUCCAUUGCUCCCACUUUUUUUUUAUUGUUUCUUUUUUUUUCCCCCUGUUGUUGUCACCAUUAUUAUGUCAUGUUAUGUCAUCGGUACUACGACAGUUGCAUUAUGAUAGUUAUUAUUUUUGUUCUUUCUUUUUUGUUCUCGUUUUGUGACGUUGAUGUUUGGUUUCAUUUUUAUUUUUGCGGUUACCUUUUUGUUUUUAUUCGCUUUUUUUUUCCUUUUUGGUUUAUUUAUUUGUACUUGGUGAAAAGGAAAAAAAACAAAAAAUAUAUAUAUAUAUAGAUACAGAAGCAGAGGAAGAGAGAAAGAGAGAAUGAAUGAAUGAAUGAAUACGAUGAAGAAGAUAUUGGAAUAGAGGUGGAAAGGGAGGGGGAAAAAAAAAAAAGAGGAAAGGGUGACGCGGUUGAGUUGACGGUAAACAAAACAGAAACGAUAAGAAAAAUAAUGCACAAAAUUCUUCCCCUUCUGUUCCUGUUUUUCGUUUCUUUGUUUGUUUUGUUUUGUUUCGUCUUCUGUUUGGUACCGCAUCACCCGUUACCAUGGCCCUCAAUAUGUCUUUAUUAUCAUUAUUAUUAUUAUUAUUAUUAUUACUAUCCGUUACUGUUAUCCUUCAUUGGCAUGAUGUUUUUCCGCCGAUAUUUCACAUCUUUUCAUUGACUCUUAUUAUUUUUCACUUCAUCCAUGCCGACUCUGCAGUACUUGUAGAAAUUUCAUUGAAACAGUAUUUUGACGAAAAGGAAGCAAGAAAGUUAAAAAUUAAUGUAAUGUAAUGCAAUGCAAUGUAAUGUAAUGUAAUAAAAAAGUAAUAACAAUGACAAACCUAAUAACUAUCAUAAUUAUAAGAGAAAUUGAAAGAAAGGACAAGGUAGGGUGGAAAAGGAAGCAAAAGAGGGAGAAGGGGGUGAAAAAAAAAAUUAAUUGUUUAAGGCUUGAGAAGGGAAAACGGCACUGUAGUAGAGAAUGAGAAGGAAUAAAAAUAAGUGCGUGAGUAAAUGGAUGAAUCAACAACUAAGUGAAUGAGCAUUUUCAUGUACACAAAAAAAAAAAUGAAAAGACGUGUUUGACUCACAAAGGGGAGGAAGAAUAGCAUGAAAGGUAAAUAUUUGUGUCAGAAUAAAAAAGAAACUAAGAAUAAAAAAGAAAUAAUAACAAUAAUAAAGUGAGGCAGAAAAUGAUGUUUCCACACCAUUGGGAUUGUUAAAUGUUGCGAUUUGGAGAGGAGGGAACGCGUGUUGACUGACGUGGUGAUGAAAAUUUUUUUGUUUUGUUUUGUUUUGUUUUGUUUUGUUUUGUUUGAGGGUCACACGUGUUCCACAACUCCUCCUUUUGUUUUAUUUUGUUUCCGCCCUCCCUCGUUCCCCCUUUCUGCGUUUCCCCUUUUUUUUUCUUUUUUUUUCUUUUUUUUUUUCUGUUUUCUGUUUUCUGUUUUGUUUUGUUUUUUUUUUUUGCAUCCCAUCGAUUUGAGAGUUAUAAAAGACGAGGAAAAGCGGAAUGUUUCUCGUGCGACGAGAGCUGGACAUGUAAAACACAAAGGGAAAUUAAGGAAGUAAAUAAAAGUAAAAAAAAAAGAAAAGAAGAAAAACGAAAAAGAAAAAAGGAAAAAGGAAAAAAAAAAGCACCCGAGUGGGUAGAGGAUAUGCUGGCAAUAGUGUGGUCAGUUUAUUUUAAAGAGGGAAUAUGUGGAGGGAAGGGAAAUUUUUUGAAUAUAAAUAUUUACUCCCACAACAUGCCGGAAAUAUAUAUUAAUAUUUGAAAAAAAAAAGAGAGAGAGAGAGAGAAAGAAAGAGAUGGAUUGGAAGGUAAGGUAAGGUAAAGAAGUAAAGUGAAUGAAGCGCGUUGAUAAUAAAUUAAGAGGAAAUAAAAAAUGUGAAGGAUUUGAAGAAGUUUUGUUGGUGCUACUUUCAAGUGAAACUAAACAAAAUAUGUGAAGGACGUAAUCAAUAUUUAUUUGUUUGUUUGAUGACGCUUCUAUCCAUCUGUCUUCCUUUUGCUUGUAAUUUCACUUUUGUAUUUUCCCUCCUUCCCCUCAUCGUUUGUUUUUUCAUACUUUUUUUACUUUUUUUUUGUGUUUGUAUGGUUGGUUGUUCCGCGUAUUGCGUUUAAAAAAAAAAAGUAUUGAAUAGCAUUGUCGUUCGUGGACCUGGCGCCCUAUUUUUUUUUGUUUUUUUUUGUUUUGUUUUACUUUAUUUUCACCUUGGAUAUGGGCAUAUGAUACAAAUAAAAUAAUAAUUAAAAAAGGGGAAGUGAGGCUUAGUGAACAAACGAAAAGGAAGUCAAAUGAAAUAAUAAUUUCUUUAUUAAAGUGGUAAAAGAACAAGAAUGACGUUAACGAUGAAAUGAGCUCCACAUAUGUCAGGUACUUGAAAUCACGUAAAAGAUAAAGAAUGAUUUAAAGGAAGAUUAGAGAAUAUGAGGAGUUAGAAAAGAGCAAGUAAGUAAGGGUAAGUAUAUUCUAAGGGAAAUGCUACUUGUACUUAUAAUUAAUGACAAUAAUAGGGAGGGCAAACCAAAUAUAUAUAAAUAUAAAUAAAGAUAUAUAUAUAUAUAUAUAUACAUAUAUUUGAUUAAAGGAGCGAGAGAAAAGGGGGGGCAGAAACGAACAAAAUAAAGUUAGUGAAGGAAAAAAGAACUGAAUAAUAAGGUGCUUUCCUUACCGCAUUUGUGAGCCACCCCUCAUCCCCACAUGUACGAGAGCAUUUUUCAUCGUUUUCGUUCCUCAACAAACUUUUGUGCGAUGAGGUGGUGGAUGAGGAGUGGCAGUGCCAAAAGUAAACAAAUGGAUCUAUAUUUCCAUUUUUGUUACUUAUGAAUGUUCACACCUUAUCUUGUUUUUUUUUAAGUUUUCAGCCUUUUUUUUUCGUUUUUUUCUUUUAAAUUGUGUGUGUGAGCUCUUUUUUUUUUUUACGCUUCCUCCCACCCCCAUUGCGGGAUCCGCUUUGCAAAGAGGAAAUAUUUUGAGGUAUGUGGAGGGCUCCCGUGUCCUCAUAUAUGUAAGCAGCGCUUUUUUUUCGUUAUGCUAAUUUGAAAAGGAAUUUUCAUGUAUGUACAUAGUUAAUAUAAAGAAAUAUAUUUAUAUAUUUAUAAUAUACAAUAUAUACAAGAAAUAUACAUACAUGUUUCUGUGACCGUCUAUCUAACUGUCUGACCAUCUAAUUGACUGACCGACUGGCUGUUUGUUUGUUGUUUUUUUUUUUUCGUUUGCGUGUAAACAUAGAUACAUCAUAAUGGAGUGGCCUCUCAUGUGUAUGUGCGGGUGCAUCCAUGUGUGCAUGUCUCGUGUUUCCGUUUAUUGUCAUUAUUAUUGUUUGUAUUCCUUGUUUCGCACCUUGUGUGUGUUUGCUUGUUCGUGUUCGCCUACUUUUUUUUUUGCUUUUUUUUUGUUUUUGUUUUUCCUCCUUCCCUUUUAUGUGAACCAUGGCUAUCAUUUCUUAUUAUUACUGACAUUAUAGCUAUUGUUAUUGUUUACUGUUGCUACUGACAUUCCUCCCUGGGUGAAUUUCAAUUAUUUGUGAUCUCCUUCUCGUGUUUUUCCCCCAUCUUCUUCAGUCUUUCCUUUUUCUUUUUUUUUGUCUGCUUCGUGUUGACGUUAGUUUUCUGUUUUUGAUGUGCGUGCAGUUUUUGUUGUUUCAACUAAAUUAAUUUUUCUGUUUUUUCCUUGAGGUCCUGCGCGCGGUGGCGAGAAAAAACAAAAGUAAAUGUCAAAUAAUGGCUGGUUUUUCGACAAGGUUAACAGAAUUCAAAAGGAAUAAACAGUUUUUAAAAGAGAAAAAAACGGUGAAAUAUAUAUAUAUAUAUAUAUAUUUAAAGCUGACAGAUAUACAAAUACGUGUGUGAUCCUGUGUGAUUGUGUGUUUGUGCUCACAUUAGAAAGUAAAUAAACAAAUUUAAUGUGCUUUUCUGCCGUCGAAACUUCCCUCCGCUCCCGUGUUUUGUCUUUGACGUGAUAUGCAGUGUGCGGAUGAGGUUGUUACCACUUUUCUUUUUGUUGUUGUUGUCGUUUCUUCGCUUUUUUUUGGGGGGGGGUUUCAGUUUUUGGCUUUUGGCUUUAUUAUUGCCGCUUAAUGGAUGGAGAAGGGAGAGGGAGGCGAUAGAAGCAAAGAAGAGUUCUGUUUGUUUGCUCGCUGCGGACUUCUGUUCACUUUGAUUUAAUAUCCGGAAAGGAGCAAAAAAAGAAAAGUAUAAUUCAUUUGACAUUUCUUGAGCAAGUCGUACAAUGAGUGAGCAUACAAAUACUUGCAGCAUCCGUUCUCUUCCCUCCUUAAUCUUUCUCAGACUCUUCCCCUGUCAUCUACUUUAUUGUUUAUUGUACUGUUCCCUUUCUAUUUUGUUUCUGUUUUCGUUUGUGUUUGUGUUAAAUCCGAUAAUCAUCAACAGCAUCUAACCGGUAUUUUGUUUUAUCCGUGGUAACGAAAAGAAAAAGAAAAAAGAGAGACAAUAAUAUAUCUCUGUUAAUGCCCACGUUAAACUAUAUAUUUUUGUAGGAAGGAGAGAAGAGGGUAUGGACCUGCUAAGGGAAUUACAUAUUUGUUCCCUUCUGUAUGAACCACACUUACCUACGGUUUUUGUUGUUUUUUUUUUGUCUUUGUUCCUCUUGUUUUGGUUUGGUUUUUUUUUUUGGUUUCGUUUCGUUUCUAUUUCCUUUUUUUCUAUUUGUUUUACGUCUUUGAUUGUCUCGUUGCGCGGGAAGAGGAAUACGAGUAAGGGGAGAGAACAGAAGAAAAGAAAAGGAGACAAGAAGAAAAAGGAAA**

**>1 (1-2000)**

**UGGCACAGAGGGUAACGAAGUAGGAAUUUUUGCCGCCAGCUGAGGUCGUUUACCUUGGGUUGGCUGUCAUGGAGAUAGGG**

**AAGAGAGAAGCAACAUCGCGUGCAAGGAAAAACGAACAGAAGAAUCGAUCCCUCCCCCCUCCAUGAGUCCUUCUUCGUUU**

**AUAAUUUCGGCUGUUGUUUUUGCUGUCACCUUUUUUUUUCUUCCUUCUCGUUCGCCACACCCUCUCCUCAACCUCCUUUA**

**CCUCCAUGGAUCUCUCGGCAGCAGUCCCCCUCCCCUCAUUCCCCUCAUGUUUUGGUGGUCGCCUUAUUGGCACCUCUUUU**

**CGUCAUUUUUUCCCCCCACUUGCUGCAGUUCAAGGCGUAUUUGUGGGGAGAGUAAAGAACAUAAGUGAAAUCUGAAAACA**

**AUAGAAAAGAGAGAUGAAUUAAUUAAUGAAUGAAUGGGAGAAAUAAGGAGGAGAGGUGGUGGAGAGCUGGGGAACUGUAA**

**UGAAGAAGAAAAAAAAGAAGAACGAGGGAAAGGUAGAUUCAGUGGGGGAGACAAAACAGUGAAGAAGAAUCUGGCGGAAA**

**GCAACAAUAAGUGUGAAAAUGAGAACGAAGGUGUGAAUGAUUCUUCAGUUUAAGAAACCACACGUGAAGAAGGACAGAAU**

**AUAUAAAUAAAUAUAUCUAUUUAUAUGUUGAUGUCUCAGAAGAAAAAAAGCAAAGGAGGGGAGAUAGAAGAGGUUUAAAG**

**GGAGGAAGGAAGACAGGCCCUGAAACCAAUAAAACAAAAUAAAUAAAUAAAGAAACGCGAAAUCAAAACUGAAAAUGCGA**

**AAAAAAAAACGAAGAACAAGAAGGGUGCAAUGAAGGAAUGAACUCCACAGAGAGAAUACCCGUUCAAACAUUCUUUUCUU**

**CGUUUUUCCCCUCCUCCUCUCCCCGACGCAACCGCCCCCCUUUUUUUUCUAUAAUUGGCUCUUCUGACCCACACCCCCUC**

**ACUUUCCUUCUCAAAGAACGGCAUCUUGUUUUCCCACCCUUUAUUAUUGUUAAUUUCUUUUUUCGCUUUACUAUUAUUAU**

**UAUUAUUUACCACCCCAUCACCGUUACUUUUAGUUGUUUUUUUUUUUUUUAUGCCUCUCUCCUUUCAUUUUUUUGCGCUU**

**UUGCUCCUCCUUAUUGGUUAUGUAGAGCGAUUUAUAUAAAUCUAAUAUAUAUAUAUAUAUAUAUAUUUGUAUUCAUAUAU**

**UUUAUGUCUUAUAUACAUAACUACUUUCGCGCAGAAAAGGGAAAGAGGAAAGGAGAAGAAGGGGGAAAAUAGAAAGCCCA**

**UGCACAUAUAUAUAUAUAUAUAUUUAUUUAUUUGUUUAUUUAUUUAUAUAUAUAUAUAUAUGUAUUCUUUGUACCUCCCU**

**UUCGUUUUUACUUUGUUUUUUGUUUUACCUUUUUUUUGUUUCUACUUUGUUUCUUGUUUUGUUUACUGUUAGUCACUCCU**

**UACUAUUUUUACCGCCUCUAUUAUUAUCACUAUUAUUAUUAUUAUUAUUACUAUUAUUAUUCCUAUUAUUAUUAUUAUUA**

**UUCCUAUUAUUAUUAUUAUUAUUAUUCCUAUUAUUAUUAUUAUUCCUAUUACUACUGCUAUUAUUAUUGUUAUUUCUGGG**

**GCUACUGCAGUGCUUUCCCUUUCUUUUUCUUUGGGUGUUCGCGUGGUUGAGAUUGUGAUGUGACUUCUGUUUGCUGACGA**

**UUGUUUGUUUCUAUUUGAUUGUGCCGGUGUCUGUUUUCCCCCACCCCCGUGAUGUCACGAAGAAAAACAAAAAUUAAAUU**

**GAAAAGUUUUUACUUCUCCUCGUCCAAAGCAAUUGCUCUCCUCCUUCCUUUUCUCCACACGCGCGCACGUACGUAGUGAA**

**CUAAUCAAAGAGUGAAGAAAUAAAAAUAAAAACACGCGUCGUGUUGUGGAUCCCUUUUUCGGAUCCAAUUUGCGCUAUUG**

**CUUUUUUUUGUUAGUAUUAUUGUUUUGUGCUUUUUUUUUUCUUUUUCUGUGUGUGUAAUAUGAUUUUGUCGCUUUUCAUU**

**>1.1 (1-618)**

**UGGCACAGAGGGUAACGAAGUAGGAAUUUUUGCCGCCAGCUGAGGUCGUUUACCUUGGGUUGGCUGUCAUGGAGAUAGGG**

**AAGAGAGAAGCAACAUCGCGUGCAAGGAAAAACGAACAGAAGAAUCGAUCCCUCCCCCCUCCAUGAGUCCUUCUUCGUUU**

**AUAAUUUCGGCUGUUGUUUUUGCUGUCACCUUUUUUUUUCUUCCUUCUCGUUCGCCACACCCUCUCCUCAACCUCCUUUA**

**CCUCCAUGGAUCUCUCGGCAGCAGUCCCCCUCCCCUCAUUCCCCUCAUGUUUUGGUGGUCGCCUUAUUGGCACCUCUUUU**

**CGUCAUUUUUUCCCCCCACUUGCUGCAGUUCAAGGCGUAUUUGUGGGGAGAGUAAAGAACAUAAGUGAAAUCUGAAAACA**

**AUAGAAAAGAGAGAUGAAUUAAUUAAUGAAUGAAUGGGAGAAAUAAGGAGGAGAGGUGGUGGAGAGCUGGGGAACUGUAA**

**UGAAGAAGAAAAAAAAGAAGAACGAGGGAAAGGUAGAUUCAGUGGGGGAGACAAAACAGUGAAGAAGAAUCUGGCGGAAA**

**GCAACAAUAAGUGUGAAAAUGAGAACGAAGGUGUGAAUGAUUCUUCAGUUUAAGAAA**

**>1.2 (598-1262)**

**GUGAUUCUUCAGUUUAAGAAACCACACGUGAAGAAGGACAGAAUAUAUAAAUAAAUAUAUCUAUUUAUAUGUUGAUGUCU**

**CAGAAGAAAAAAAGCAAAGGAGGGGAGAUAGAAGAGGUUUAAAGGGAGGAAGGAAGACAGGCCCUGAAACCAAUAAAACA**

**AAAUAAAUAAAUAAAGAAACGCGAAAUCAAAACUGAAAAUGCGAAAAAAAAAACGAAGAACAAGAAGGGUGCAAUGAAGG**

**AAUGAACUCCACAGAGAGAAUACCCGUUCAAACAUUCUUUUCUUCGUUUUUCCCCUCCUCCUCUCCCCGACGCAACCGCC**

**CCCCUUUUUUUUCUAUAAUUGGCUCUUCUGACCCACACCCCCUCACUUUCCUUCUCAAAGAACGGCAUCUUGUUUUCCCA**

**CCCUUUAUUAUUGUUAAUUUCUUUUUUCGCUUUACUAUUAUUAUUAUUAUUUACCACCCCAUCACCGUUACUUUUAGUUG**

**UUUUUUUUUUUUUUAUGCCUCUCUCCUUUCAUUUUUUUGCGCUUUUGCUCCUCCUUAUUGGUUAUGUAGAGCGAUUUAUA**

**UAAAUCUAAUAUAUAUAUAUAUAUAUAUAUUUGUAUUCAUAUAUUUUAUGUCUUAUAUACAUAACUACUUUCGCGCAGAA**

**AAGGGAAAGAGGAAAGGAGAAGAAGG**

**>1.2 ΔAU**

**GUGAUUCUUCAGUUUAAGAAACCACACGUGAAGAAGGACAGAAUAUAUAAAUAAAUAUAUCUAUUUAUAUGUUGAUGUCUCAGAAGAAAAAAAGCAAAGGAGGGGAGAUAGAAGAGGUUUAAAGGGAGGAAGGAAGACAGGCCCUGAAACCAAUAAAACAAAAUAAAUAAAUAAAGAAACGCGAAAUCAAAACUGAAAAUGCGAAAAAAAAAACGAAGAACAAGAAGGGUGCAAUGAAGGAAUGAACUCCACAGAGAGAAUACCCGUUCAAACAUUCUUUUCUUCGUUUUUCCCCUCCUCCUCUCCCCGACGCAACCGCCCCCCUUUUUUUUCUAUAAUUGGCUCUUCUGACCCACACCCCCUCACUUUCCUUCUCAAAGAACGGCAUCUUGUUUUCCCACCCUUUAUUAUUGUUAAUUUCUUUUUUCGCUUUACUAUUAUUAUUAUUAUUUACCACCCCAUCACCGUUACUUUUAGUUGUUUUUUUUUUUUUUAUGCCUCUCUCCUUUCAUUUUUUUGCGCUUUUGCUCCUCCUUAUUGGUUAUGUAGAGCGAUUUAUAUAAAUCUAA......................UUUGUAUUCAUAUAUUUUAUGUCUUAUAUACAUAACUACUUUCGCGCAGAAAAGGGAAAGAGGAAAGGAGAAGAAGG**

**> 1.3 (1239-1772)**

**GGGAAAGAGGAAAGGAGAAGAAGGGGGAAAAUAGAAAGCCCAUGCACAUAUAUAUAUAUAUAUAUUUAUUUAUUUGUUUA**

**UUUAUUUAUAUAUAUAUAUAUAUGUAUUCUUUGUACCUCCCUUUCGUUUUUACUUUGUUUUUUGUUUUACCUUUUUUUUG**

**UUUCUACUUUGUUUCUUGUUUUGUUUACUGUUAGUCACUCCUUACUAUUUUUACCGCCUCUAUUAUUAUCACUAUUAUUA**

**UUAUUAUUAUUACUAUUAUUAUUCCUAUUAUUAUUAUUAUUAUUCCUAUUAUUAUUAUUAUUAUUAUUCCUAUUAUUAUU**

**AUUAUUCCUAUUACUACUGCUAUUAUUAUUGUUAUUUCUGGGGCUACUGCAGUGCUUUCCCUUUCUUUUUCUUUGGGUGU**

**UCGCGUGGUUGAGAUUGUGAUGUGACUUCUGUUUGCUGACGAUUGUUUGUUUCUAUUUGAUUGUGCCGGUGUCUGUUUUC**

**CCCCACCCCCGUGAUGUCACGAAGAAAAACAAAAAUUAAAUUGAAAAGUUUUUA**

**> 1.4 (1766-2000)**

**GUUUUUACUUCUCCUCGUCCAAAGCAAUUGCUCUCCUCCUUCCUUUUCUCCACACGCGCGCACGUACGUAGUGAACUAAU**

**CAAAGAGUGAAGAAAUAAAAAUAAAAACACGCGUCGUGUUGUGGAUCCCUUUUUCGGAUCCAAUUUGCGCUAUUGCUUUU**

**UUUUGUUAGUAUUAUUGUUUUGUGCUUUUUUUUUUCUUUUUCUGUGUGUGUAAUAUGAUUUUGUCGCUUUUCAUU**

**> 1.4.1 (1766-1879)**

**GUUUUUACUUCUCCUCGUCCAAAGCAAUUGCUCUCCUCCUUCCUUUUCUCCACACGCGCGCACGUACGUAGUGAACUAAU**

**CAAAGAGUGAAGAAAUAAAAAUAAAAACACGCG**

**> 1.4.2 (1874-2000)**

**CACGCGUCGUGUUGUGGAUCCCUUUUUCGGAUCCAAUUUGCGCUAUUGCUUUUUUUUGUUAGUAUUAUUGUUUUGUGCUUUUUUUUUUCUUUUUCUGUGUGUGUAAUAUGAUUUUGUCGCUUUUCAUU**

**>2 (2001-4023)**

**CAGCGGAAGCAGACAGUAAAGUUAUGGGCUCAUAUGCUUGCAUGUGCAUUCCAUCCGAUAUUGCGCGGGAGUUGUUGUUU**

**UGUUUUGUUUUUCACACCUUCGUCCCUUUUUUUUUUCUUUUUGGUUUUUAUUGUUUGUAUAUAUUUUCAUAUUUAUGUAC**

**AUAUAUUAGGAGCAAAUGCAUGCGUGUUUGUGCUUUAACGCCCAUGUGUGCGCCCCUUGGCUUGCUUUCCCACAAAUACC**

**CUUUCGAAACGCCUUACGGAAGCGGGGACAAUAUAAUUAAGAAGAAAGGGAGAUGGGGGGGAAAAAAAAAAGAGAAAGAG**

**AAAGAGAAACACCGGCUCGUAGACAUGAGGAGAGGAAUCAAAAAAAAGACAAUGAAAUGAAAUAAAGUUAAGUGAAGUGA**

**AUUGAAACGAGGUAAGAAUUUAAGAAGUUAAGAAAGCUGGUGACCCGCCGUCGCUUUUCUACUUCAUUCCACUCUUUUUU**

**UUUCUUCUUUUCUUUUUUUUCCCAUCCUUUUCCUUUUUUUUUGCUAGUUUUUGAUCCGCUUUGCCUUUGCAUCUUAUCUU**

**UCCAUUGCUCCCACUUUUUUUUUAUUGUUUCUUUUUUUUUCCCCCUGUUGUUGUCACCAUUAUUAUGUCAUGUUAUGUCA**

**UCGGUACUACGACAGUUGCAUUAUGAUAGUUAUUAUUUUUGUUCUUUCUUUUUUGUUCUCGUUUUGUGACGUUGAUGUUU**

**GGUUUCAUUUUUAUUUUUGCGGUUACCUUUUUGUUUUUAUUCGCUUUUUUUUUCCUUUUUGGUUUAUUUAUUUGUACUUG**

**GUGAAAAGGAAAAAAAACAAAAAAUUAUAUAUAUAUAGAUACAGAAGCAGAGGAAGAGAGAAAGAGAGAAUGAAUGAAUG**

**AAUGAAUACGAUGAAGAAGAUAUUGGAAUAGAGGUGGAAAGGGAGGGGGAAAAAAAAAAAAGAGGAAAGGGUGACGCGGU**

**UGAGUUGACGGUAAACAAAACAGAAACGAUAAGAAAAAUAAUGCACAAAAUUCUUCCCCUUCUGUUCCUGUUUUUCGUUU**

**CUUUGUUUGUUUUGUUUUGUUUCGUCUUCUGUUUGGUACCGCAUCACCCGUUACCAUGGCCCUCAAUAUGUCUUUAUUAU**

**CAUUAUUAUUAUUAUUAUUAUUAUUACUAUCCGUUACUGUUAUCCUUCAUUGGCAUGAUGUUUUUCCGCCGAUAUUUCAC**

**AUCUUUUCAUUGACUCUUAUUAUUUUUCACUUCAUCCAUGCCGACUCUGCAGUACUUGUAGAAAUUUCAUUGAAACAGUA**

**UUUUGACGAAAAGGAAGCAAGAAAGUUAAAAAUUAAUGUAAUGUAAUGCAAUGCAAUGUAAUGUAAUGUAAUAAAAAAGU**

**AAUAACAAUGACAAACCUAAUAACUAUCAUAAUUAUAAGAGAAAUUGAAAGAAAGGACAAGGUAGGGUGGAAAAGGAAGC**

**AAAAGAGGGAGAAGGGGGUGAAAAAAAAAAUUAAUUGUUUAAGGCUUGAGAAGGGAAAACGGCACUGUAGUAGAGAAUGA**

**GAAGGAAUAAAAAUAAGUGCGUGAGUAAAUGGAUGAAUCAACAACUAAGUGAAUGAGCAUUUUCAUGUACACAAAAAAAA**

**AAAUGAAAAGACGUGUUUGACUCACAAAGGGGAGGAAGAAUAGCAUGAAAGGUAAAUAUUUGUGUCAGAAUAAAAAAGAA**

**ACUAAGAAUAAAAAAGAAAUAAUAACAAUAAUAAAGUGAGGCAGAAAAUGAUGUUUCCACACCAUUGGGAUUGUUAAAUG**

**UUGCGAUUUGGAGAGGAGGGAACGCGUGUUGACUGACGUGGUGAUGAAAAUUUUUUUGUUUUGUUUUGUUUUGUUUUGUU**

**UUGUUUUGUUUGAGGGUCACACGUGUUCCACAACUCCUCCUUUUGUUUUAUUUUGUUUCCGCCCUCCCUCGUUCCCCCUU**

**UCUGCGUUUCCCCUUUUUUUUUCUUUUUUUUUCUUUUUUUUUUUCUGUUUUCUGUUUUCUGUUUUGUUUUGUUUUUUUUU**

**UUUGCAUCCCAUCGAUUUGAGAG**

**>2.1 (2001-2552)**

**CAGCGGAAGCAGACAGUAAAGUUAUGGGCUCAUAUGCUUGCAUGUGCAUUCCAUCCGAUAUUGCGCGGGAGUUGUUGUUU**

**UGUUUUGUUUUUCACACCUUCGUCCCUUUUUUUUUUCUUUUUGGUUUUUAUUGUUUGUAUAUAUUUUCAUAUUUAUGUAC**

**AUAUAUUAGGAGCAAAUGCAUGCGUGUUUGUGCUUUAACGCCCAUGUGUGCGCCCCUUGGCUUGCUUUCCCACAAAUACC**

**CUUUCGAAACGCCUUACGGAAGCGGGGACAAUAUAAUUAAGAAGAAAGGGAGAUGGGGGGGAAAAAAAAAAGAGAAAGAG**

**AAAGAGAAACACCGGCUCGUAGACAUGAGGAGAGGAAUCAAAAAAAAGACAAUGAAAUGAAAUAAAGUUAAGUGAAGUGA**

**AUUGAAACGAGGUAAGAAUUUAAGAAGUUAAGAAAGCUGGUGACCCGCCGUCGCUUUUCUACUUCAUUCCACUCUUUUUU**

**UUUCUUCUUUUCUUUUUUUUCCCAUCCUUUUCCUUUUUUUUUGCUAGUUUUUGAUCCGCUUUGCCUUUGCAU**

**>2.2 (2533-4023)**

**GAUCCGCUUUGCCUUUGCAUCUUAUCUUUCCAUUGCUCCCACUUUUUUUUUAUUGUUUCUUUUUUUUUCCCCCUGUUGUU**

**GUCACCAUUAUUAUGUCAUGUUAUGUCAUCGGUACUACGACAGUUGCAUUAUGAUAGUUAUUAUUUUUGUUCUUUCUUUU**

**UUGUUCUCGUUUUGUGACGUUGAUGUUUGGUUUCAUUUUUAUUUUUGCGGUUACCUUUUUGUUUUUAUUCGCUUUUUUUU**

**UCCUUUUUGGUUUAUUUAUUUGUACUUGGUGAAAAGGAAAAAAAACAAAAAAUUAUAUAUAUAUAGAUACAGAAGCAGAG**

**GAAGAGAGAAAGAGAGAAUGAAUGAAUGAAUGAAUACGAUGAAGAAGAUAUUGGAAUAGAGGUGGAAAGGGAGGGGGAAA**

**AAAAAAAAAGAGGAAAGGGUGACGCGGUUGAGUUGACGGUAAACAAAACAGAAACGAUAAGAAAAAUAAUGCACAAAAUU**

**CUUCCCCUUCUGUUCCUGUUUUUCGUUUCUUUGUUUGUUUUGUUUUGUUUCGUCUUCUGUUUGGUACCGCAUCACCCGUU**

**ACCAUGGCCCUCAAUAUGUCUUUAUUAUCAUUAUUAUUAUUAUUAUUAUUAUUACUAUCCGUUACUGUUAUCCUUCAUUG**

**GCAUGAUGUUUUUCCGCCGAUAUUUCACAUCUUUUCAUUGACUCUUAUUAUUUUUCACUUCAUCCAUGCCGACUCUGCAG**

**UACUUGUAGAAAUUUCAUUGAAACAGUAUUUUGACGAAAAGGAAGCAAGAAAGUUAAAAAUUAAUGUAAUGUAAUGCAAU**

**GCAAUGUAAUGUAAUGUAAUAAAAAAGUAAUAACAAUGACAAACCUAAUAACUAUCAUAAUUAUAAGAGAAAUUGAAAGA**

**AAGGACAAGGUAGGGUGGAAAAGGAAGCAAAAGAGGGAGAAGGGGGUGAAAAAAAAAAUUAAUUGUUUAAGGCUUGAGAA**

**GGGAAAACGGCACUGUAGUAGAGAAUGAGAAGGAAUAAAAAUAAGUGCGUGAGUAAAUGGAUGAAUCAACAACUAAGUGA**

**AUGAGCAUUUUCAUGUACACAAAAAAAAAAAUGAAAAGACGUGUUUGACUCACAAAGGGGAGGAAGAAUAGCAUGAAAGG**

**UAAAUAUUUGUGUCAGAAUAAAAAAGAAACUAAGAAUAAAAAAGAAAUAAUAACAAUAAUAAAGUGAGGCAGAAAAUGAU**

**GUUUCCACACCAUUGGGAUUGUUAAAUGUUGCGAUUUGGAGAGGAGGGAACGCGUGUUGACUGACGUGGUGAUGAAAAUU**

**UUUUUGUUUUGUUUUGUUUUGUUUUGUUUUGUUUUGUUUGAGGGUCACACGUGUUCCACAACUCCUCCUUUUGUUUUAUU**

**UUGUUUCCGCCCUCCCUCGUUCCCCCUUUCUGCGUUUCCCCUUUUUUUUUCUUUUUUUUUCUUUUUUUUUUUCUGUUUUC**

**UGUUUUCUGUUUUGUUUUGUUUUUUUUUUUUGCAUCCCAUCGAUUUGAGAG**

**> 2.2.1 (2533-3229)**

**GAUCCGCUUUGCCUUUGCAUCUUAUCUUUCCAUUGCUCCCACUUUUUUUUUAUUGUUUCUUUUUUUUUCCCCCUGUUGUU**

**GUCACCAUUAUUAUGUCAUGUUAUGUCAUCGGUACUACGACAGUUGCAUUAUGAUAGUUAUUAUUUUUGUUCUUUCUUUU**

**UUGUUCUCGUUUUGUGACGUUGAUGUUUGGUUUCAUUUUUAUUUUUGCGGUUACCUUUUUGUUUUUAUUCGCUUUUUUUU**

**UCCUUUUUGGUUUAUUUAUUUGUACUUGGUGAAAAGGAAAAAAAACAAAAAAUUAUAUAUAUAUAGAUACAGAAGCAGAG**

**GAAGAGAGAAAGAGAGAAUGAAUGAAUGAAUGAAUACGAUGAAGAAGAUAUUGGAAUAGAGGUGGAAAGGGAGGGGGAAA**

**AAAAAAAAAGAGGAAAGGGUGACGCGGUUGAGUUGACGGUAAACAAAACAGAAACGAUAAGAAAAAUAAUGCACAAAAUU**

**CUUCCCCUUCUGUUCCUGUUUUUCGUUUCUUUGUUUGUUUUGUUUUGUUUCGUCUUCUGUUUGGUACCGCAUCACCCGUU**

**ACCAUGGCCCUCAAUAUGUCUUUAUUAUCAUUAUUAUUAUUAUUAUUAUUAUUACUAUCCGUUACUGUUAUCCUUCAUUG**

**GCAUGAUGUUUUUCCGCCGAUAUUUCACAUCUUUUCAUUGACUCUUAUUAUUUUUCA**

**> 2.2.2 (3230-4023)**

**CUUCAUCCAUGCCGACUCUGCAGUACUUGUAGAAAUUUCAUUGAAACAGUAUUUUGACGAAAAGGAAGCAAGAAAGUUAA**

**AAAUUAAUGUAAUGUAAUGCAAUGCAAUGUAAUGUAAUGUAAUAAAAAAGUAAUAACAAUGACAAACCUAAUAACUAUCA**

**UAAUUAUAAGAGAAAUUGAAAGAAAGGACAAGGUAGGGUGGAAAAGGAAGCAAAAGAGGGAGAAGGGGGUGAAAAAAAAA**

**AUUAAUUGUUUAAGGCUUGAGAAGGGAAAACGGCACUGUAGUAGAGAAUGAGAAGGAAUAAAAAUAAGUGCGUGAGUAAA**

**UGGAUGAAUCAACAACUAAGUGAAUGAGCAUUUUCAUGUACACAAAAAAAAAAAUGAAAAGACGUGUUUGACUCACAAAG**

**GGGAGGAAGAAUAGCAUGAAAGGUAAAUAUUUGUGUCAGAAUAAAAAAGAAACUAAGAAUAAAAAAGAAAUAAUAACAAU**

**AAUAAAGUGAGGCAGAAAAUGAUGUUUCCACACCAUUGGGAUUGUUAAAUGUUGCGAUUUGGAGAGGAGGGAACGCGUGU**

**UGACUGACGUGGUGAUGAAAAUUUUUUUGUUUUGUUUUGUUUUGUUUUGUUUUGUUUUGUUUGAGGGUCACACGUGUUCC**

**ACAACUCCUCCUUUUGUUUUAUUUUGUUUCCGCCCUCCCUCGUUCCCCCUUUCUGCGUUUCCCCUUUUUUUUUCUUUUUU**

**UUUCUUUUUUUUUUUCUGUUUUCUGUUUUCUGUUUUGUUUUGUUUUUUUUUUUUGCAUCCCAUCGAUUUGAGAG**

**>2.2.3 (3230-3730)**

**CUUCAUCCAUGCCGACUCUGCAGUACUUGUAGAAAUUUCAUUGAAACAGUAUUUUGACGAAAAGGAAGCAAGAAAGUUAA**

**AAAUUAAUGUAAUGUAAUGCAAUGCAAUGUAAUGUAAUGUAAUAAAAAAGUAAUAACAAUGACAAACCUAAUAACUAUCA**

**UAAUUAUAAGAGAAAUUGAAAGAAAGGACAAGGUAGGGUGGAAAAGGAAGCAAAAGAGGGAGAAGGGGGUGAAAAAAAAA**

**AUUAAUUGUUUAAGGCUUGAGAAGGGAAAACGGCACUGUAGUAGAGAAUGAGAAGGAAUAAAAAUAAGUGCGUGAGUAAA**

**UGGAUGAAUCAACAACUAAGUGAAUGAGCAUUUUCAUGUACACAAAAAAAAAAAUGAAAAGACGUGUUUGACUCACAAAG**

**GGGAGGAAGAAUAGCAUGAAAGGUAAAUAUUUGUGUCAGAAUAAAAAAGAAACUAAGAAUAAAAAAGAAAUAAUAACAAU**

**AAUAAAGUGAGGCAGAAAAUG**

**> 2.2.4 (3731-4023)**

**AUGUUUCCACACCAUUGGGAUUGUUAAAUGUUGCGAUUUGGAGAGGAGGGAACGCGUGUUGACUGACGUGGUGAUGAAAA**

**UUUUUUUGUUUUGUUUUGUUUUGUUUUGUUUUGUUUUGUUUGAGGGUCACACGUGUUCCACAACUCCUCCUUUUGUUUUA**

**UUUUGUUUCCGCCCUCCCUCGUUCCCCCUUUCUGCGUUUCCCCUUUUUUUUUCUUUUUUUUUCUUUUUUUUUUUCUGUUU**

**UCUGUUUUCUGUUUUGUUUUGUUUUUUUUUUUUGCAUCCCAUCGAUUUGAGAG**

**>2.2.5 (3731-3866)**

**AUGUUUCCACACCAUUGGGAUUGUUAAAUGUUGCGAUUUGGAGAGGAGGGAACGCGUGUUGACUGACGUGGUGAUGAAAAUUUUUUUGUUUUGUUUUGUUUUGUUUUGUUUUGUUUUGUUUGAGGGUCACACGUG**

**>2.2.6 (3859-4023)**

**CACGUGUUCCACAACUCCUCCUUUUGUUUUAUUUUGUUUCCGCCCUCCCUCGUUCCCCCUUUCUGCGUUUCCCCUUUUUUUUUCUUUUUUUUUCUUUUUUUUUUUCUGUUUUCUGUUUUCUGUUUUGUUUUGUUUUUUUUUUUUGCAUCCCAUCGAUUUGAGAG**

**>3 (4024-5999)**

**UUAUAAAAGACGAGGAAAAGCGGAAUGUUUCUCGUGCGACGAGAGCUGGACAUGUAAAACACAAAGGGAAAUUAAGGAAG**

**UAAAUAAAAGUAAAAAAAAAAGAAAAGAAGAAAAACGAAAAAGAAAAAAGGAAAAAGGAAAAAAAAAAGCACCCGAGUGG**

**GUAGAGGAUAUGCUGGCAAUAGUGUGGUCAGUUUAUUUUAAAGAGGGAAUAUGUGGAGGGAAGGGAAAUUUUUUGAAUAU**

**AAAUAUUUACUCCCACAACAUGCCGGAAAUAUAUAUUAAUAUUUGAAAAAAAAAAGAGAGAGAGAGAGAGAAAGAAAGAG**

**AUGGAUUGGAAGGUAAGGUAAGGUAAAGAAGUAAAGUGAAUGAAGCGCGUUGAUAAUAAAUUAAGAGGAAAUAAAAAAUG**

**UGAAGGAUUUGAAGAAGUUUUGUUGGUGCUACUUUCAAGUGAAACUAAACAAAAUAUGUGAAGGACGUAAUCAAUAUUUA**

**UUUGUUUGUUUGAUGACGCUUCUAUCCAUCUGUCUUCCUUUUGCUUGUAAUUUCACUUUUGUAUUUUCCCUCCUUCCCCU**

**CAUCGUUUGUUUUUUCAUACUUUUUUUACUUUUUUUUUGUGUUUGUAUGGUUGGUUGUUCCGCGUAUUGCGUUUAAAAAA**

**AAAAAGUAUUGAAUAGCAUUGUCGUUCGUGGACCUGGCGCCCUAUUUUUUUUUGUUUUUUUUUGUUUUGUUUUACUUUAU**

**UUUCACCUUGGAUAUGGGCAUAUGAUACAAAUAAAAUAAUAAUUAAAAAAGGGGAAGUGAGGCUUAGUGAACAAACGAAA**

**AGGAAGUCAAAUGAAAUAAUAAUUUCUUUAUUAAAGUGGUAAAAGAACAAGAAUGACGUUAACGAUGAAAUGAGCUCCAC**

**AUAUGUCAGGUACUUGAAAUCACGUAAAAGAUAAAGAAUGAUUUAAAGGAAGAUUAGAGAAUAUGAGGAGUUAGAAAAGA**

**GCAAGUAAGUAAGGGUAAGUAUAUUCUAAGGGAAAUGCUACUUGUACUUAUAAUUAAUGACAAUAAUAGGGAGGGCAAAC**

**CAAAUAUAUAUAAAUAUAAAUAAAGAUAUAUAUAUAUAUAUAUAUACAUAUAUUUGAUUAAAGGAGCGAGAGAAAAGGGG**

**GGGCAGAAACGAACAAAAUAAAGUUAGUGAAGGAAAAAAGAACUGAAUAAUAAGGUGCUUUCCUUACCGCAUUUGUGAGC**

**CACCCCUCAUCCCCACAUGUACGAGAGCAUUUUUCAUCGUUUUCGUUCCUCAACAAACUUUUGUGCGAUGAGGUGGUGGA**

**UGAGGAGUGGCAGUGCCAAAAGUAAACAAAUGGAUCUAUAUUUCCAUUUUUGUUACUUAUGAAUGUUCACACCUUAUCUU**

**GUUUUUUUUUAAGUUUUCAGCCUUUUUUUUUCGUUUUUUUCUUUUAAAUUGUGUGUGUGAGCUCUUUUUUUUUUUUACGC**

**UUCCUCCCACCCCCAUUGCGGGAUCCGCUUUGCAAAGAGGAAAUAUUUUGAGGUAUGUGGAGGGCUCCCGUGUCCUCAUA**

**UAUGUAAGCAGCGCUUUUUUUUCGUUAUGCUAAUUUGAAAAGGAAUUUUCAUGUAUGUACAUAGUUAAUAUAAAGAAAUA**

**UAUUUAUAUAUUUAUAAUAUACAAUAUAUACAAGAAAUAUACAUACAUGUUUCUGUGACCGUCUAUCUAACUGUCUGACC**

**AUCUAAUUGACUGACCGACUGGCUGUUUGUUUGUUGUUUUUUUUUUUUCGUUUGCGUGUAAACAUAGAUACAUCAUAAUG**

**GAGUGGCCUCUCAUGUGUAUGUGCGGGUGCAUCCAUGUGUGCAUGUCUCGUGUUUCCGUUUAUUGUCAUUAUUAUUGUUU**

**GUAUUCCUUGUUUCGCACCUUGUGUGUGUUUGCUUGUUCGUGUUCGCCUACUUUUUUUUUUGCUUUUUUUUUGUUUUUGU**

**UUUUCCUCCUUCCCUUUUAUGUGAACCAUGGCUAUCAUUUCUUAUUAUUACUGACA**

**> 3.1 (4024-4531)**

**UUAUAAAAGACGAGGAAAAGCGGAAUGUUUCUCGUGCGACGAGAGCUGGACAUGUAAAACACAAAGGGAAAUUAAGGAAG**

**UAAAUAAAAGUAAAAAAAAAAGAAAAGAAGAAAAACGAAAAAGAAAAAAGGAAAAAGGAAAAAAAAAAGCACCCGAGUGG**

**GUAGAGGAUAUGCUGGCAAUAGUGUGGUCAGUUUAUUUUAAAGAGGGAAUAUGUGGAGGGAAGGGAAAUUUUUUGAAUAU**

**AAAUAUUUACUCCCACAACAUGCCGGAAAUAUAUAUUAAUAUUUGAAAAAAAAAAGAGAGAGAGAGAGAGAAAGAAAGAG**

**AUGGAUUGGAAGGUAAGGUAAGGUAAAGAAGUAAAGUGAAUGAAGCGCGUUGAUAAUAAAUUAAGAGGAAAUAAAAAAUG**

**UGAAGGAUUUGAAGAAGUUUUGUUGGUGCUACUUUCAAGUGAAACUAAACAAAAUAUGUGAAGGACGUAAUCAAUAUUUA**

**UUUGUUUGUUUGAUGACGCUUCUAUCCA**

**>3.1.2 (4139-4531)**

**CGAAAAAGAAAAAAGGAAAAAGGAAAAAAAAAAGCACCCGAGUGGGUAGAGGAUAUGCUGGCAAUAGUGUGGUCAGUUUAUUUUAAAGAGGGAAUAUGUGGAGGGAAGGGAAAUUUUUUGAAUAUAAAUAUUUACUCCCACAACAUGCCGGAAAUAUAUAUUAAUAUUUGAAAAAAAAAAGAGAGAGAGAGAGAGAAAGAAAGAGAUGGAUUGGAAGGUAAGGUAAGGUAAAGAAGUAAAGUGAAUGAAGCGCGUUGAUAAUAAAUUAAGAGGAAAUAAAAAAUGUGAAGGAUUUGAAGAAGUUUUGUUGGUGCUACUUUCAAGUGAAACUAAACAAAAUAUGUGAAGGACGUAAUCAAUAUUUAUUUGUUUGUUUGAUGACGCUUCUAUCCA**

**>3.1.2 ΔAAGG(A_10_)**

**CGAAAAAGAAAAAAGGAAA..............GCACCCGAGUGGGUAGAGGAUAUGCUGGCAAUAGUGUGGUCAGUUUAUUUUAAAGAGGGAAUAUGUGGAGGGAAGGGAAAUUUUUUGAAUAUAAAUAUUUACUCCCACAACAUGCCGGAAAUAUAUAUUAAUAUUUGAAAAAAAAAAGAGAGAGAGAGAGAGAAAGAAAGAGAUGGAUUGGAAGGUAAGGUAAGGUAAAGAAGUAAAGUGAAUGAAGCGCGUUGAUAAUAAAUUAAGAGGAAAUAAAAAAUGUGAAGGAUUUGAAGAAGUUUUGUUGGUGCUACUUUCAAGUGAAACUAAACAAAAUAUGUGAAGGACGUAAUCAAUAUUUAUUUGUUUGUUUGAUGACGCUUCUAUCCAG**

**>3.2 (4520-5999)**

**CGCUUCUAUCCAUCUGUCUUCCUUUUGCUUGUAAUUUCACUUUUGUAUUUUCCCUCCUUCCCCUCAUCGUUUGUUUUUUC**

**AUACUUUUUUUACUUUUUUUUUGUGUUUGUAUGGUUGGUUGUUCCGCGUAUUGCGUUUAAAAAAAAAAAGUAUUGAAUAG**

**CAUUGUCGUUCGUGGACCUGGCGCCCUAUUUUUUUUUGUUUUUUUUUGUUUUGUUUUACUUUAUUUUCACCUUGGAUAUG**

**GGCAUAUGAUACAAAUAAAAUAAUAAUUAAAAAAGGGGAAGUGAGGCUUAGUGAACAAACGAAAAGGAAGUCAAAUGAAA**

**UAAUAAUUUCUUUAUUAAAGUGGUAAAAGAACAAGAAUGACGUUAACGAUGAAAUGAGCUCCACAUAUGUCAGGUACUUG**

**AAAUCACGUAAAAGAUAAAGAAUGAUUUAAAGGAAGAUUAGAGAAUAUGAGGAGUUAGAAAAGAGCAAGUAAGUAAGGGU**

**AAGUAUAUUCUAAGGGAAAUGCUACUUGUACUUAUAAUUAAUGACAAUAAUAGGGAGGGCAAACCAAAUAUAUAUAAAUA**

**UAAAUAAAGAUAUAUAUAUAUAUAUAUAUACAUAUAUUUGAUUAAAGGAGCGAGAGAAAAGGGGGGGCAGAAACGAACAA**

**AAUAAAGUUAGUGAAGGAAAAAAGAACUGAAUAAUAAGGUGCUUUCCUUACCGCAUUUGUGAGCCACCCCUCAUCCCCAC**

**AUGUACGAGAGCAUUUUUCAUCGUUUUCGUUCCUCAACAAACUUUUGUGCGAUGAGGUGGUGGAUGAGGAGUGGCAGUGC**

**CAAAAGUAAACAAAUGGAUCUAUAUUUCCAUUUUUGUUACUUAUGAAUGUUCACACCUUAUCUUGUUUUUUUUUAAGUUU**

**UCAGCCUUUUUUUUUCGUUUUUUUCUUUUAAAUUGUGUGUGUGAGCUCUUUUUUUUUUUUACGCUUCCUCCCACCCCCAU**

**UGCGGGAUCCGCUUUGCAAAGAGGAAAUAUUUUGAGGUAUGUGGAGGGCUCCCGUGUCCUCAUAUAUGUAAGCAGCGCUU**

**UUUUUUCGUUAUGCUAAUUUGAAAAGGAAUUUUCAUGUAUGUACAUAGUUAAUAUAAAGAAAUAUAUUUAUAUAUUUAUA**

**AUAUACAAUAUAUACAAGAAAUAUACAUACAUGUUUCUGUGACCGUCUAUCUAACUGUCUGACCAUCUAAUUGACUGACC**

**GACUGGCUGUUUGUUUGUUGUUUUUUUUUUUUCGUUUGCGUGUAAACAUAGAUACAUCAUAAUGGAGUGGCCUCUCAUGU**

**GUAUGUGCGGGUGCAUCCAUGUGUGCAUGUCUCGUGUUUCCGUUUAUUGUCAUUAUUAUUGUUUGUAUUCCUUGUUUCGC**

**ACCUUGUGUGUGUUUGCUUGUUCGUGUUCGCCUACUUUUUUUUUUGCUUUUUUUUUGUUUUUGUUUUUCCUCCUUCCCUU**

**UUAUGUGAACCAUGGCUAUCAUUUCUUAUUAUUACUGACA**

**>3.2.1 (4520-5214)**

**CGCUUCUAUCCAUCUGUCUUCCUUUUGCUUGUAAUUUCACUUUUGUAUUUUCCCUCCUUCCCCUCAUCGUUUGUUUUUUC**

**AUACUUUUUUUACUUUUUUUUUGUGUUUGUAUGGUUGGUUGUUCCGCGUAUUGCGUUUAAAAAAAAAAAGUAUUGAAUAG**

**CAUUGUCGUUCGUGGACCUGGCGCCCUAUUUUUUUUUGUUUUUUUUUGUUUUGUUUUACUUUAUUUUCACCUUGGAUAUG**

**GGCAUAUGAUACAAAUAAAAUAAUAAUUAAAAAAGGGGAAGUGAGGCUUAGUGAACAAACGAAAAGGAAGUCAAAUGAAA**

**UAAUAAUUUCUUUAUUAAAGUGGUAAAAGAACAAGAAUGACGUUAACGAUGAAAUGAGCUCCACAUAUGUCAGGUACUUG**

**AAAUCACGUAAAAGAUAAAGAAUGAUUUAAAGGAAGAUUAGAGAAUAUGAGGAGUUAGAAAAGAGCAAGUAAGUAAGGGU**

**AAGUAUAUUCUAAGGGAAAUGCUACUUGUACUUAUAAUUAAUGACAAUAAUAGGGAGGGCAAACCAAAUAUAUAUAAAUA**

**UAAAUAAAGAUAUAUAUAUAUAUAUAUAUACAUAUAUUUGAUUAAAGGAGCGAGAGAAAAGGGGGGGCAGAAACGAACAA**

**AAUAAAGUUAGUGAAGGAAAAAAGAACUGAAUAAUAAGGUGCUUUCCUUACCGCA**

**>3.2.2 (5208-5999)**

**UACCGCAUUUGUGAGCCACCCCUCAUCCCCACAUGUACGAGAGCAUUUUUCAUCGUUUUCGUUCCUCAACAAACUUUUGU**

**GCGAUGAGGUGGUGGAUGAGGAGUGGCAGUGCCAAAAGUAAACAAAUGGAUCUAUAUUUCCAUUUUUGUUACUUAUGAAU**

**GUUCACACCUUAUCUUGUUUUUUUUUAAGUUUUCAGCCUUUUUUUUUCGUUUUUUUCUUUUAAAUUGUGUGUGUGAGCUC**

**UUUUUUUUUUUUACGCUUCCUCCCACCCCCAUUGCGGGAUCCGCUUUGCAAAGAGGAAAUAUUUUGAGGUAUGUGGAGGG**

**CUCCCGUGUCCUCAUAUAUGUAAGCAGCGCUUUUUUUUCGUUAUGCUAAUUUGAAAAGGAAUUUUCAUGUAUGUACAUAG**

**UUAAUAUAAAGAAAUAUAUUUAUAUAUUUAUAAUAUACAAUAUAUACAAGAAAUAUACAUACAUGUUUCUGUGACCGUCU**

**AUCUAACUGUCUGACCAUCUAAUUGACUGACCGACUGGCUGUUUGUUUGUUGUUUUUUUUUUUUCGUUUGCGUGUAAACA**

**UAGAUACAUCAUAAUGGAGUGGCCUCUCAUGUGUAUGUGCGGGUGCAUCCAUGUGUGCAUGUCUCGUGUUUCCGUUUAUU**

**GUCAUUAUUAUUGUUUGUAUUCCUUGUUUCGCACCUUGUGUGUGUUUGCUUGUUCGUGUUCGCCUACUUUUUUUUUUGCU**

**UUUUUUUUGUUUUUGUUUUUCCUCCUUCCCUUUUAUGUGAACCAUGGCUAUCAUUUCUUAUUAUUACUGACA**

**>4 (6000-8081)**

**AAAAGGAAA: possible polyadenylation site**

**AG: possible splice acceptor site for downstream pseudogene RNA**

**Lower case - possible intergenic region**

**UUAUAGCUAUUGUUAUUGUUUACUGUUGCUACUGACAUUCCUCCCUGGGUGAAUUUCAAUUAUUUGUGAUCUCCUUCUCG**

**UGUUUUUCCCCCAUCUUCUUCAGUCUUUCCUUUUUCUUUUUUUUUGUCUGCUUCGUGUUGACGUUAGUUUUCUGUUUUUG**

**AUGUGCGUGCAGUUUUUGUUGUUUCAACUAAAUUAAUUUUUCUGUUUUUUCCUUGAGGUCCUGCGCGCGGUGGCGAGAAA**

**AAACAAAAGUAAAUGUCAAAUAAUGGCUGGUUUUUCGACAAGGUUAACAGAAUUCAAAAGGAAUAAACAGUUUUUAAAAG**

**AGAAAAAAACGGUGAAAUAUAUAUAUAUAUAUAUAUAUUUAAAGCUGACAGAUAUACAAAUACGUGUGUGAUCCUGUGUG**

**AUUGUGUGUUUGUGCUCACAUUAGAAAGUAAAUAAACAAAUUUAAUGUGCUUUUCUGCCGUCGAAACUUCCCUCCGCUCC**

**CGUGUUUUGUCUUUGACGUGAUAUGCAGUGUGCGGAUGAGGUUGUUACCACUUUUCUUUUUGUUGUUGUUGUCGUUUCUU**

**CGCUUUUUUUUGGGGGGGGGUUUCAGUUUUUGGCUUUUGGCUUUAUUAUUGCCGCUUAAUGGAUGGAGAAGGGAGAGGGA**

**GGCGAUAGAAGCAAAGAAGAGUUCUGUUUGUUUGCUCGCUGCGGACUUCUGUUCACUUUGAUUUAAUAUCCGGAAAGGAG**

**CAAAAAAAGAAAAGUAUAAUUCAUUUGACAUUUCUUGAGCAAGUCGUACAAUGAGUGAGCAUACAAAUACUUGCAGCAUC**

**CGUUCUCUUCCCUCCUUAAUCUUUCUCAGACUCUUCCCCUGUCAUCUACUUUAUUGUUUAUUGUACUGUUCCCUUUCUAU**

**UUUGUUUCUGUUUUCGUUUGUGUUUGUGUUAAAUCCGAUAAUCAUCAACAGCAUCUAACCGGUAUUUUGUUUUAUCCGUG**

**GUAACGAAAAGAAAAAGAAAAAAGAGAGACAAUAAUAUAUCUCUGUUAAUGCCCACGUUAAACUAUAUAUUUUUGUAGGA**

**AGGAGAGAAGAGGGUAUGGACCUGCUAAGGGAAUUACAUAUUUGUUCCCUUCUGUAUGAACCACACUUACCUACGGUUUU**

**UGUUGUUUUUUUUUUGUCUUUGUUCCUCUUGUUUUGGUUUGGUUUUUUUUUUUGGUUUCGUUUCGUUUCUAUUUCCUUUU**

**UUUCUAUUUGUUUUACGUCUUUGAUUGUCUCGUUGCGCGGGAAGAGGAAUACGAGUAAGGGGAGAGAACAGAAGAAAAGA**

**AAAGGAGACAAGAAGAAAAAGGAAAuuuuguuuauuccuuaugcuuucaaaacucuuuucccguaucagugugcguguac**

**gugugugacugccucgauacaacugguuucacuuucgugcccuaucaucugcuccuucucuuuaccucuuuuuuuuucuu**

**uuuucuguuccuuucauucugcucucacguucaacaaccacacAGUGAAUUUUAUUAUUUAUUAACUUUAUUAUUACGAC**

**AAUAUAUAUAUAUACGGUCGCUACUGUUAGCUUACCGUCAAUAUCAGUAACCAUGUCACGCGGUUCACAAAUGUUAAAUU**

**UAAUAACAGUAAUAAAACGAAUAUACAAUUGAAUAGACGUGCAGAAGUUUCGAAGUUAAGUCGAAAGAGCAAAAAAGUUA**

**AAGAAAAAAAUAUGUAGAGGGUGGAGCCAACAAUAACGUAUGGGAGGAAUGAAUGUGCUUUAUGUCCGUGACGGAGGCGU**

**AAAAUGAUGAUGAACUGAAGUUAUGGAAACUAACAAAAUAAGGGAAAGGACAUAUAUAUAUAUAUAUAUAUGCGUGCAUA**

**UAUUGUGCCGGAAGGAAGCGGAAAUAUACACUGAAGCGUACCGGUCAAAAAUUGAAAGUGCGGAACCCCGCGUCUCCGCG**

**UAGUGUCACCUGUUAUCCGCGCAUUUUCUUAGUGUCUCUGCUUUUAGCGUUGCACAUCACCUUUGCAACAACCGAGUUAU**

**CGCCGUUUCACACCUGUGGCACAGAUUUUCGUGGCUCCUUCAUGACACUUAUGCUCACUUUACAUCUCACCUGCCGUUUU**

**U**

**>4.1 (6000-6699)**

**UUAUAGCUAUUGUUAUUGUUUACUGUUGCUACUGACAUUCCUCCCUGGGUGAAUUUCAAUUAUUUGUGAUCUCCUUCUCG**

**UGUUUUUCCCCCAUCUUCUUCAGUCUUUCCUUUUUCUUUUUUUUUGUCUGCUUCGUGUUGACGUUAGUUUUCUGUUUUUG**

**AUGUGCGUGCAGUUUUUGUUGUUUCAACUAAAUUAAUUUUUCUGUUUUUUCCUUGAGGUCCUGCGCGCGGUGGCGAGAAA**

**AAACAAAAGUAAAUGUCAAAUAAUGGCUGGUUUUUCGACAAGGUUAACAGAAUUCAAAAGGAAUAAACAGUUUUUAAAAG**

**AGAAAAAAACGGUGAAAUAUAUAUAUAUAUAUAUAUAUUUAAAGCUGACAGAUAUACAAAUACGUGUGUGAUCCUGUGUG**

**AUUGUGUGUUUGUGCUCACAUUAGAAAGUAAAUAAACAAAUUUAAUGUGCUUUUCUGCCGUCGAAACUUCCCUCCGCUCC**

**CGUGUUUUGUCUUUGACGUGAUAUGCAGUGUGCGGAUGAGGUUGUUACCACUUUUCUUUUUGUUGUUGUUGUCGUUUCUU**

**CGCUUUUUUUUGGGGGGGGGUUUCAGUUUUUGGCUUUUGGCUUUAUUAUUGCCGCUUAAUGGAUGGAGAAGGGAGAGGGA**

**GGCGAUAGAAGCAAAGAAGAGUUCUGUUUGUUUGCUCGCUGCGGACUUCUGUUCACUUUG**

**>4.2 (6681-7301)**

**GCGGACUUCUGUUCACUUUGAUUUAAUAUCCGGAAAGGAGCAAAAAAAGAAAAGUAUAAUUCAUUUGACAUUUCUUGAGC**

**AAGUCGUACAAUGAGUGAGCAUACAAAUACUUGCAGCAUCCGUUCUCUUCCCUCCUUAAUCUUUCUCAGACUCUUCCCCU**

**GUCAUCUACUUUAUUGUUUAUUGUACUGUUCCCUUUCUAUUUUGUUUCUGUUUUCGUUUGUGUUUGUGUUAAAUCCGAUA**

**AUCAUCAACAGCAUCUAACCGGUAUUUUGUUUUAUCCGUGGUAACGAAAAGAAAAAGAAAAAAGAGAGACAAUAAUAUAU**

**CUCUGUUAAUGCCCACGUUAAACUAUAUAUUUUUGUAGGAAGGAGAGAAGAGGGUAUGGACCUGCUAAGGGAAUUACAUA**

**UUUGUUCCCUUCUGUAUGAACCACACUUACCUACGGUUUUUGUUGUUUUUUUUUUGUCUUUGUUCCUCUUGUUUUGGUUU**

**GGUUUUUUUUUUUGGUUUCGUUUCGUUUCUAUUUCCUUUUUUUCUAUUUGUUUUACGUCUUUGAUUGUCUCGUUGCGCGG**

**GAAGAGGAAUACGAGUAAGGGGAGAGAACAGAAGAAAAGAAAAGGAGACAAGAAGAAAAAG**

**>ACT**

**UAACACCGGGUUGUGUGGCCAAAAUUGUUCUGUAGUCGCUGUGAGUUGACACGGCUAGUGCUUAUGAUUUUCCUCGCGUGUGGUGCCUGUACUCAGCCCUAUGCCUUAUUUGCAACACAUUUACGUACAGCGCACAAGAGAAGAGAAGAUCACUUGAAGAUAAUAAAUAUAGGGUUGUAGGCAUCUUGUUUAACUCAAAUUUUCUCGUCUUGGUGU**
